# The evolutionary fate of Neanderthal DNA in 30,780 admixed genomes with recent African-like ancestry

**DOI:** 10.1101/2024.07.25.605203

**Authors:** Aaron Pfennig, Joseph Lachance

## Abstract

Following introgression, Neanderthal DNA was initially purged from non-African genomes, but the evolutionary fate of remaining introgressed DNA has not been explored yet. To fill this gap, we analyzed 30,780 admixed genomes with African-like ancestry from the All of Us research program, in which Neanderthal alleles encountered novel genetic backgrounds during the last 15 generations. Observed amounts of Neanderthal DNA approximately match expectations based on ancestry proportions, suggesting neutral evolution. Nevertheless, we identified genomic regions that have significantly less or more Neanderthal ancestry than expected and are associated with spermatogenesis, innate immunity, and other biological processes. We also identified three novel introgression desert-like regions in recently admixed genomes, whose genetic features are compatible with hybrid incompatibilities and intrinsic negative selection. Overall, we find that much of the remaining Neanderthal DNA in human genomes is not under strong selection, and complex evolutionary dynamics have shaped introgression landscapes in our species.

## 1 Introduction

The sequencing of the Neanderthal genome revealed that modern humans interbred with archaic hominins after the out-of-Africa migration ~50 thousand years ago (kya) (Green et al., 2010; Prüfer et al., 2017), leaving present-day non-Africans with ~1-2% Neanderthal ancestry (Sankararaman et al., 2014; Vernot and Akey, 2014; Sankararaman et al., 2016; Vernot et al., 2016; Skov et al., 2020; Witt et al., 2023). However, the initial introgression pulse was likely greater than 5% (Harris and Nielsen, 2016; Iasi et al., 2024), indicating that much of the Neanderthal DNA was purged from modern human genomes. This purging occurred quickly as the amount of Neanderthal ancestry remained constant for the last 45,000 years in Europe (Petr et al., 2019; Iasi et al., 2024). Observations that this purging was particularly pronounced from functional genomic elements (Dannemann et al., 2017; Telis et al., 2020) and that archaic haplotypes do not carry more deleterious variants than non-archaic haplotypes in present-day Icelandic genomes (Skov et al., 2020) suggest that remaining Neanderthal DNA in extant genomes is evolutionary neutral. However, the evolutionary fate of Neanderthal DNA in contemporary populations has yet to be assessed at biobank scale.

A striking feature of the introgression landscapes in Eurasian populations are large introgression deserts, i.e., genomic regions ≥8 Mb significantly depleted of archaic introgression (Sankararaman et al., 2014; Vernot and Akey, 2014; Vernot et al., 2016; Sankararaman et al., 2016). However, the evolutionary mechanisms behind the introgression deserts are still debated. While some studies invoked hybrid incompatibilities as an explanation (Sankararaman et al., 2014, 2016; Harris et al., 2023), others argued that intrinsic negative selection against Neanderthal alleles due to their higher mutational load is a more parsimonious explanation for introgression deserts (Juric et al., 2016; Vernot et al., 2016; Harris and Nielsen, 2016; Kim et al., 2018; Steinrücken et al., 2018; Petr et al., 2019). From a theoretical population genetic perspective, both explanations are plausible (Uecker et al., 2015; Sachdeva and Barton, 2018a,b; Pfennig and Lachance, 2022).

Here, we leverage whole-genome sequences of 30,780 recently admixed individuals with pre-dominantly African-like and European-like ancestry from the United States in All of Us (All of Us Research Program Investigators et al., 2019; Bick et al., 2024) to directly test the evolutionary fate of remaining Neanderthal segments in extant human genomes. Because African genomes contain no or only very little Neanderthal ancestry (Chen et al., 2020), many archaic haplotypes have only been exposed to an African genetic background during the last 15 generations (Figure 1). This novel genetic context offers a unique opportunity to infer the evolutionary impact of Neanderthal DNA. Assuming neutrality of the remaining archaic variants, the Neanderthal introgression landscape in such admixed genomes only depends on the introgression landscape in the admixing populations and recent ancestry patterns. Thus, observing less or more Neanderthal introgressed sequence than expected based on ancestry patterns can be indicative of recent negative or positive secondary selection in these admixed genomes, respectively. Furthermore, admixed genomes with African-like ancestry potentially allow the evolutionary dynamics behind introgression deserts to be interrogated. We note that recent selection of Neanderthal DNA in admixed genomes has not yet been exhaustively tested, although a recent study by Witt et al. (2023) described the introgression landscape in admixed populations in the Americas and identified several candidates for adaptive introgression using the population branch statistic.

**Fig. 1.**
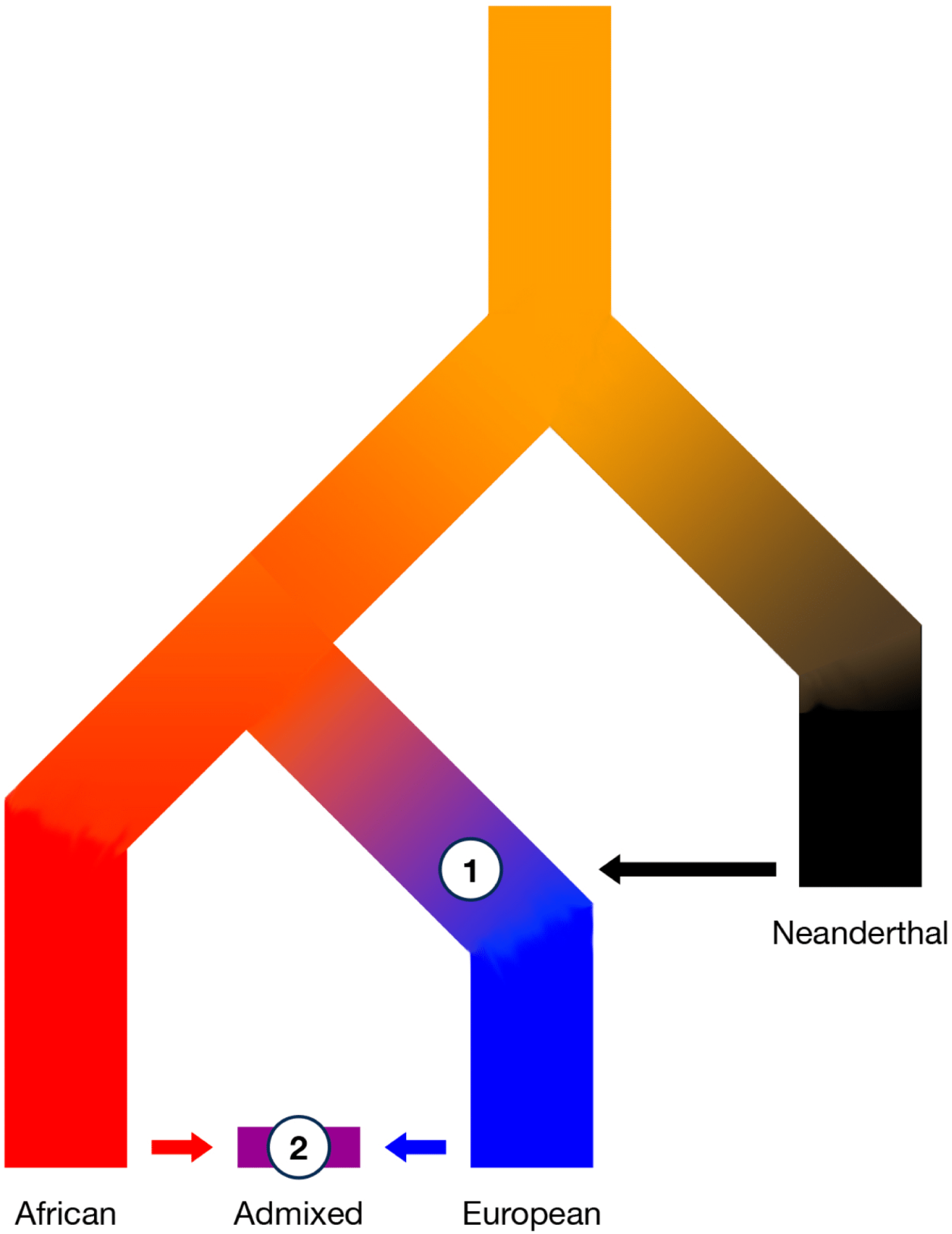
Secondary contact has brought Neanderthal DNA into novel genomic contexts. (1) Neanderthal DNA introgressed into non-African populations ~50 kya, leading to an initial purging of Neanderthal ancestry. (2) During the past 15 generations, recent admixture of individuals with African-like ancestry and European-like ancestry has introduced Neanderthal variants into a novel genetic background, potentially leading to secondary selection.

We first test the evolutionary fate of remaining Neanderthal DNA on a genome level by modeling the expected amount of introgressed sequence in these admixed genomes based on recent ancestry proportions and average amounts of introgressed sequence in the respective continental reference populations. Subsequently, we extend this model to individual genomic regions and identify potential target loci of secondary selection in the admixed individuals. Lastly, we provide new insights into the evolutionary dynamics of archaic introgression deserts by interrogating novel desert-like regions in these recently admixed genomes.

## 2 Results

We identified 30,780 recently, mostly two-way admixed individuals with predominantly African-like and European-like ancestry in All of Us, using previously inferred ancestry proportions (All of Us Research Program Investigators et al., 2019; Conley et al., 2023; Bick et al., 2024). To ensure that Neanderthal DNA was introduced into novel genetic backgrounds, i.e., African-like ancestry, we only included admixed individuals with at least 50% African-like ancestry, at least 10% European-like ancestry, and at least 95% African-like + European-like ancestry in our study. On average, the analyzed admixed individuals have 80.1% African-like, 18.3% European-like, and 1.6% East Asian/-Native American-like ancestry (Figure S1). Furthermore, we constructed continental reference panels of Neanderthal introgression landscapes using unadmixed individuals with African-like (1,067 individuals), European-like (10,503), and East Asian/Native American-like (575 individuals) ancestry from the 1000 genomes project (1KGP) (Auton et al., 2015) and All of Us (All of Us Research Program Investigators et al., 2019; Bick et al., 2024). Due to the paucity of Native American-like reference genomes and since they have previously been shown to have similar amounts of Neanderthal introgressed sequence as East Asian genomes (Sankararaman et al., 2016), we pooled East Asian and Native American genomes into one panel.

### 2.1 Inference of Neanderthal introgressed segments in global populations

Using IBDmix and the Vindija33.19 Neanderthal reference genome (Chen et al., 2020; Prüfer et al., 2017), we separately identified Neanderthal introgressed segments in the recently admixed individuals and each continental reference subpopulation and used the Denisovan reference genome to control for incomplete lineage sorting (ILS) (see Materials and Methods). We refer to this call set of Neanderthal introgressed segments as the “unfiltered” call set. Note that we only considered autosomal data. Individuals with East Asian/Native American-like and European-like ancestry show the highest amount of Neanderthal ancestry with, on average, 54.2 and 48.7 Mb per individual, respectively, while individuals with African-like ancestry have, on average, 12.9 Mb putatively Neanderthal introgressed sequence per individual. Admixed genomes with recent African-like and European-like ancestry contain intermediate amounts of Neanderthal ancestry, i.e., on average, 23.1 Mb per individual (Figure 2A). The amounts of Neanderthal ancestry in the admixed genomes are negatively correlated with recent African-like ancestry and positively correlated with recent European-like ancestry (Figure 2B - C). Due to our sampling scheme they are only weakly correlated with the recent East Asian/Native American-like ancestry (Figure 2D). Furthermore, predicted Neanderthal segments in admixed genomes are also enriched in regions with recent European-like ancestry, as opposed to African-like ancestry (Figure S3).

**Fig. 2.**
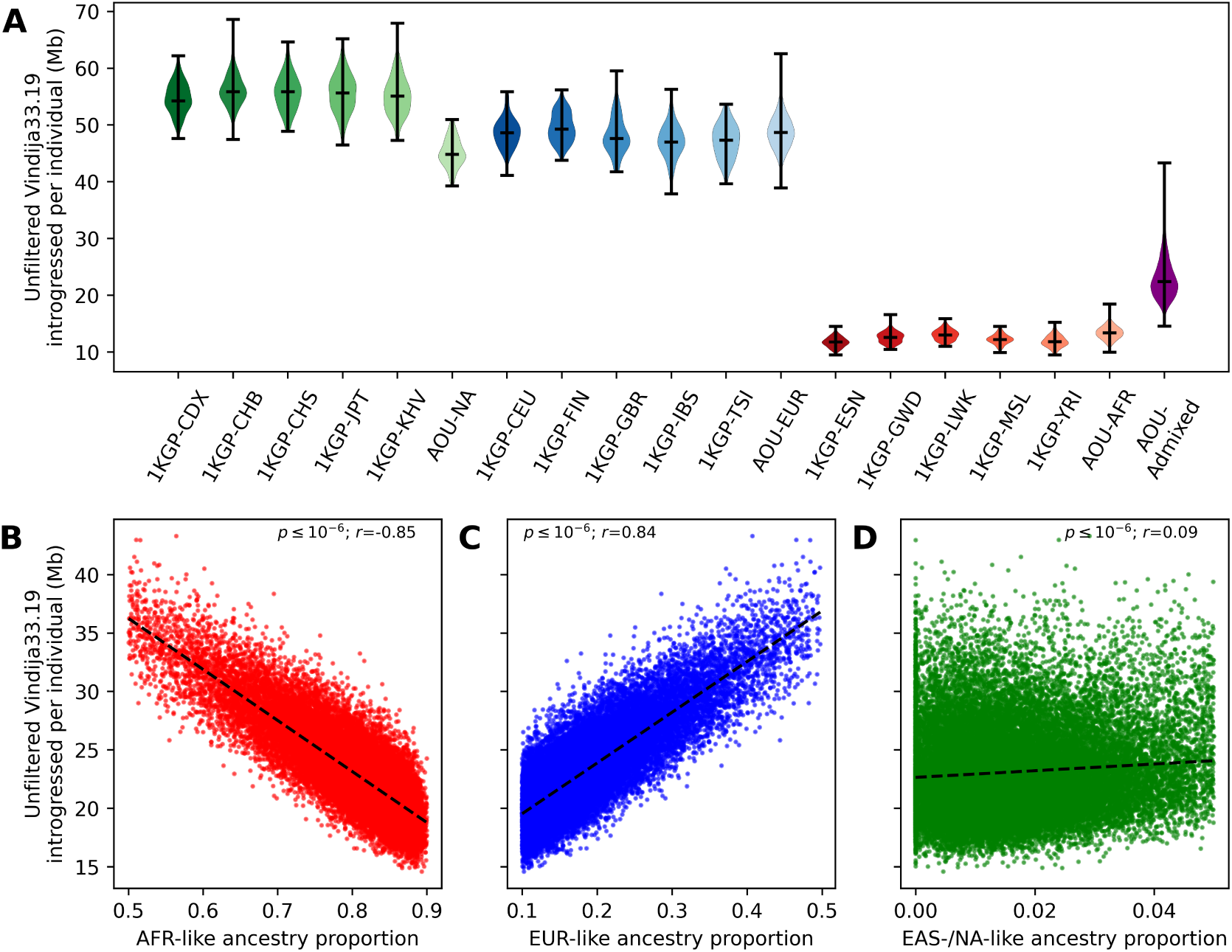
Amounts of Neanderthal ancestry in global populations and correlations with recent ancestry proportions in 30,780 admixed individuals from All of Us. **A)** Inferred amounts of Neanderthal ancestry in Mb per individual for different continental reference subpopulations, using IBDmix. East Asian/Native American populations (green) show the highest amounts of Neanderthal ancestry, immediately followed by European populations (blue). African populations (red) have the lowest amounts of inferred Neanderthal ancestry. Admixed genomes (purple; AOU-Admixed) contain intermediate amounts of Neanderthal ancestry. The whiskers indicate 1.5 times the inter-quartile range. See also Figure S2 for amounts of introgressed sequence per individual after applying the African mask **B)** The amount of Neanderthal ancestry in admixed genomes is negatively correlated with the African-like (AFR-like) ancestry proportion and **C)** positively correlated with the European-like (EUR-like) ancestry proportion. **D)** Due to our inclusion criteria, there is only a weak correlation between the amount of Neanderthal ancestry and the amount of East Asian/-Native American-like (EAS/NA-like) ancestry in the admixed genomes. The p-value (*p*) and Pearson’s correlation coefficient (*r*) for separate linear regressions are given in the respective panels. See also Figure S3.

### 2.2 No evidence for polygenic selection of Neanderthal ancestry on a genome level since admixture

Using the introgression landscape in African, European, and East Asian/Native American reference populations and estimated ancestry proportions, we modeled the expected amounts of Neanderthal introgressed sequence in recently admixed genomes as a linear mixture of the continental reference populations (Equation 1, see Materials and Methods). If Neanderthal ancestry is effectively neutral in extant genomes, as indirectly suggested by previous studies (Harris and Nielsen, 2016; Petr et al., 2019; Skov et al., 2020; Wei et al., 2023), one would observe as much Neanderthal introgressed sequence as expected based on recent ancestry patterns in the admixed genomes and average amounts of introgressed sequence from continental reference populations. Whereas, if Neanderthal ancestry is selected against or for, one would expect to see less or more Neanderthal ancestry in recently admixed genomes than expected, respectively. Within European-like and African-like continental ancestry groups, individuals from different populations show similar amounts of inferred Neanderthal introgressed sequence (e.g., compare AOU-EUR and 1KGP-EUR reference populations in Figure 2A), suggesting little confounding in our modeling from continental heterogeneity in the admixing African and European populations and not knowing the exact genetic ancestry of the admixing populations 15 generations ago. By contrast, individuals with Native American-like ancestry from All of Us (AOU-NA) have less introgressed sequence than East Asian 1KGP reference populations (Figure 2A), although a previous study found that they have similar amounts of Neanderthal DNA (Sankararaman et al., 2016). However, potential differences in the introgression landscapes between East Asian and Native American populations should also not bias subsequent analyses as we limited our analysis to individuals with less than 5% recent East Asian/Native American-like ancestry and there is only a weak correlation of Neanderthal introgression amounts and recent East Asian/Native American-like ancestry proportions in the admixed individuals analyzed here (Figure 2D).

We found that expected and observed amounts of Neanderthal ancestry per individual are strongly correlated (*p* ≤ 10^−6^; Pearson’s correlation *r* = 0.85). Despite this pattern, we observed more Neanderthal ancestry in the recently admixed genomes than expected (Figure S4A). However, we also observed this pattern in neutral coalescent simulations under a plausible demographic model (Figure S4B). Although this enrichment is robust to variation in recombination rate (Figure S5), we show below that this enrichment is the result of ILS and false positive predictions.

To account for remaining biases from ILS and false positive predictions, we removed any introgressed segment that overlapped with a predicted segment in African reference genomes for all subsequent analyses. This was done for two reasons. First, despite including Argweaver-D predicted human-to-Neanderthal introgressed regions in the mask for IBDmix (Hubisz et al., 2020) and using the Denisovan reference genome to control for ILS (see Materials and Methods), IBD-mix still predicts Neanderthal introgressed segments with a higher “false-positive” rate in African genomes due to earlier human-to-Neanderthal introgression events (Harris et al., 2023; Li et al., 2024). Indeed, introgressed segments removed using this “African mask” show characteristics of false positive predictions. They are shorter (Mann-Whitney U *p* ≤ 10^−6^; Figure S6A), have lower LOD scores (Mann-Whitney U *p* ≤ 10^−6^; Figure S6B), and are in regions with lower recombination rates (Mann-Whitney U *p* ≤ 10^−6^; Figure S6C). Second, regardless of whether introgressed segments in African reference genomes are true or false positive predictions, we are only interested in the evolutionary dynamics of Neanderthal haplotypes that were not present in an African genetic background before admixture 15 generations ago. Only the fitness of these Neanderthal haplotypes has been truly re-assessed in the admixed genomes. After removing Neanderthal segments overlapping with introgressed segments in African reference genomes, European and East Asian/Native American reference genomes contain, on average, 12.6 Mb and 21.7 Mb introgressed Neanderthal sequence, respectively, and the average amount of Neanderthal ancestry per admixed genome is reduced to, on average, 3.0 Mb (Figure S2).

When focusing on introgressed segments that were mostly contributed by European ancestors and modeling expected amounts of introgressed sequence per individual based on this African masked call set (Equation 1), we still observe a strong correlation between expected and observed amounts of Neanderthal introgressed sequence per admixed individual (*p* ≤ 10^−6^; *r* = 0.79). However, we observed only slightly more Neanderthal ancestry than expected in the admixed individuals (Figure 3A). The differences between expected and observed admixture fractions are significantly reduced and centered near zero with a mean difference of 0.34 Mb (0.012% of the entire genome). Analyzing simulated data in the same way removed the initially observed Neanderthal enrichment (Figure 3B), indicating that previously observed biases from ILS and false positive predictions are corrected by applying the African mask. Thus, despite the initially observed enrichment, there is no evidence for strong, polygenic selection of Neanderthal introgressed segments that were newly introduced into an African genetic background 15 generations ago on a genome level in admixed individuals with recent African-like and European-like ancestry. We replicated these results on a smaller test dataset consisting of 93 admixed individuals from 1KGP-ACB and 1KGP-ASW as well as using the other two available high-quality Neanderthal reference genomes (i.e., Altai and Chagyrskaya) (see Supplemental Information, Figure S7).

**Fig. 3.**
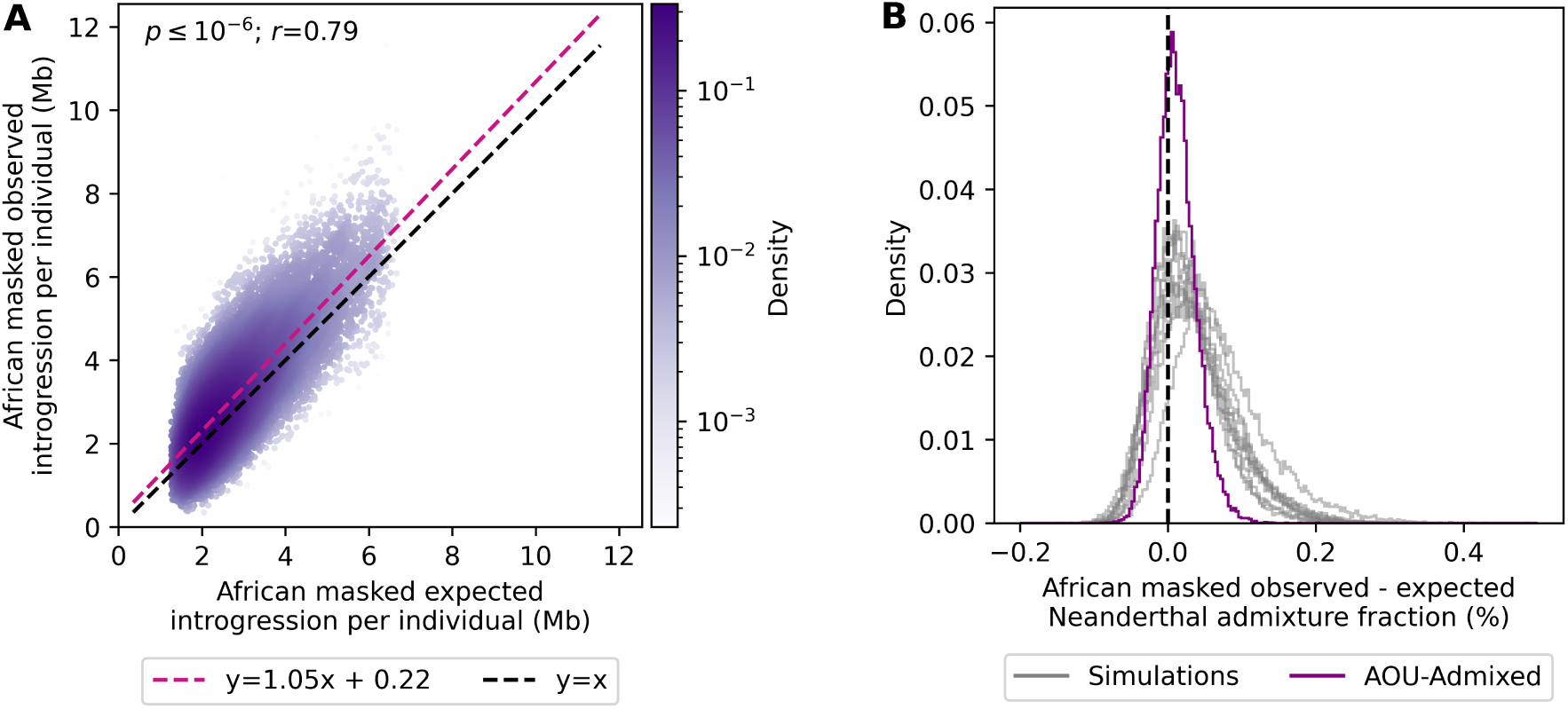
Observed amounts of Neanderthal ancestry per individual are largely compatible with neutral evolution in 30,780 admixed genomes from All of Us after correcting for incomplete lineage sorting and false positives by removing segments that overlapped with putative Neanderthal segments in African reference genomes. **A)** Slightly more Neanderthal ancestry is observed than expected, but the slope of the regression line is close to one (m=1.05, 95%CI: 1.04 - 1.05), and the y-intercept is close to zero (b=0.22, 95% CI:0.19 - 0.24). The p-value (*p*) and Pearson’s correlation coefficient (*r*) of the regression line are given in the panel. **B)** Differences in expected and observed Neanderthal admixture fractions are centered near zero for empirical data (purple) and data from neutral coalescence simulations (gray). The mean difference in the Neanderthal admixture fraction in the empirical data is 0.34 Mb (0.012% of the entire genome). See also Figures S4, S5, S6, and S7.

### 2.3 Regions with significantly less or more Neanderthal ancestry than expected affect known Neanderthal phenotypes

Despite not finding evidence for strong, polygenic selection of remaining Neanderthal ancestry on a genome level, individual regions may still be under selection. To identify regions with significantly less or more Neanderthal ancestry than expected, we first painted local ancestry using FLARE (Browning et al., 2023), i.e., we identified whether genomic segments in admixed genomes had recent African-like, European-like, or East Asian/Native American-like ancestry. We calculated local ancestry and Neanderthal introgression frequencies for overlapping 50 kb windows (10 kb strides). Using these local ancestry and introgression frequencies from our African masked call set, we modeled the expected number of Neanderthal haplotypes as independent binomial draws from all reference populations, i.e., a multinomial distribution (Equation 2; see Materials and Methods). As before, on a genome level, we observe a strong correlation between expected and observed introgression frequencies (*p* ≤ 10^−6^; *r* = 0.94; Figure S8A), and the differences between expected and observed introgression frequencies are centered near zero, indicating that our modeling approach is not inherently biased (Figure S8B).

Using neutral simulations, we found that probabilistic modeling under the above-described model was not well calibrated to identify windows with significantly less or more Neanderthal ancestry than expected (Equation 3 and Equation S1 in Supplemental Information), and in particular, regions with significantly more Neanderthal ancestry than expected appeared to be false positives (Figure S9). To be more conservative in identifying outliers and accounting for genetic drift, we, therefore, conditioned our analysis of 50 kb genomic windows on the aggregated results of neutral coalescence simulations (Figure 4A & B). We subtracted the simulated joint spectrum of expected and observed introgression frequencies from the empirical joint spectrum and searched for peaks in the residual spectrum, using the Watershed algorithm (see Materials and Methods). That is, we identified regions with a higher density of windows with specific expected and observed introgression frequencies in the empirical spectrum than could be expected under neutral evolution and a plausible demographic model. Using this approach, we identified two peaks in the empirical joint spectrum with windows depleted and enriched for Neanderthal ancestry relative to expectations, respectively (ellipses in Figure 4C). These windows formed four and three independent genomic regions with significantly less and more Neanderthal ancestry than expected, respectively (Figure 5; Table S1). Notably, all of these regions were also identified using probabilistic modeling assuming binomial inheritance (Equation 3 and Equation S1 in Supplemental Information).

**Fig. 4.**
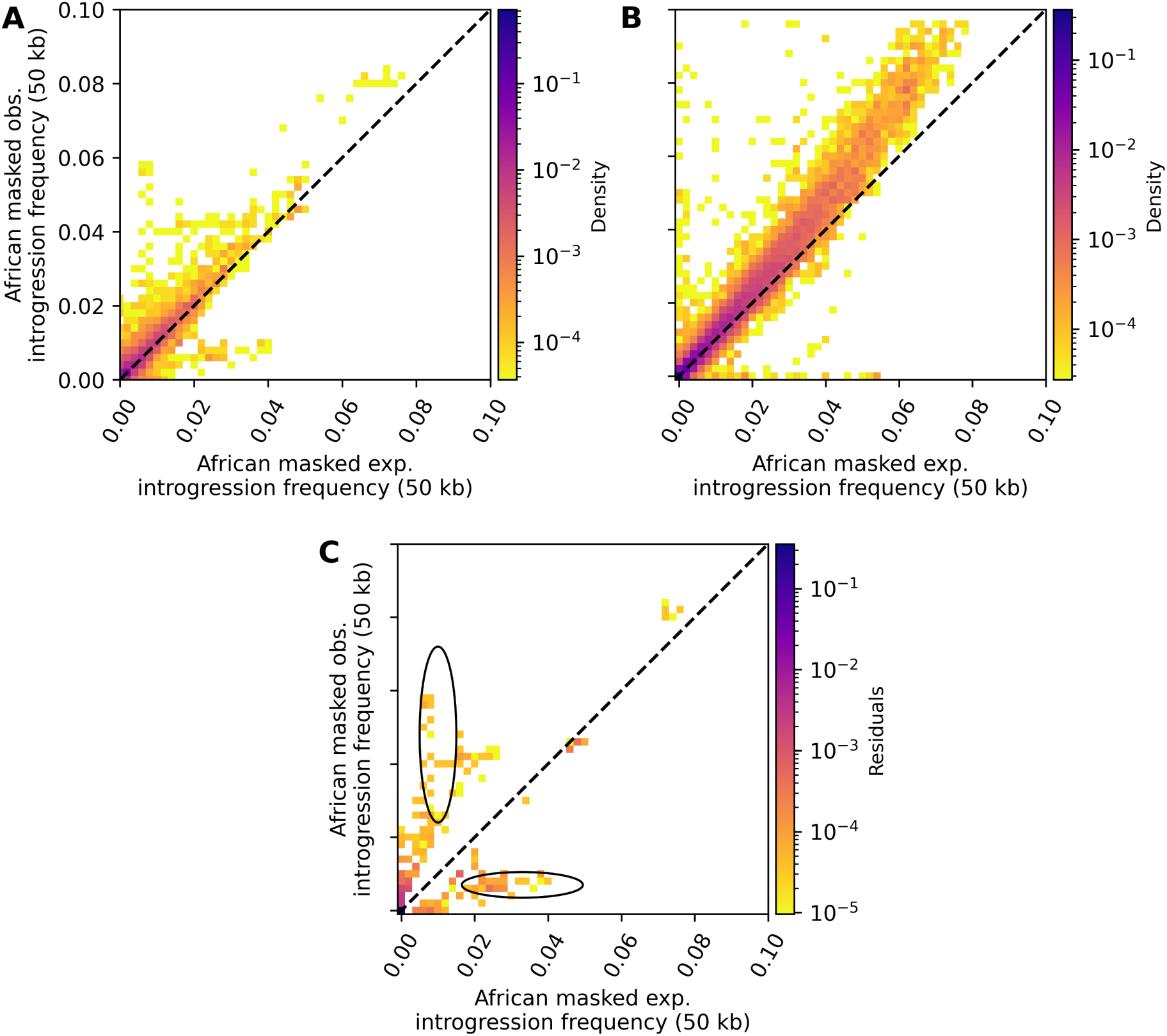
Spectra of expected vs. observed Neanderthal introgression frequencies in 50 kb windows after applying the African mask for empirical and simulated data. That is, we removed any Neanderthal segment that overlapped with a predicted segment in African reference genomes. **A)** and **B)** show expected vs observed introgression frequencies in the 30,780 admixed individuals from All of Us and aggregated simulated data, respectively. **C)** shows the positive residuals when panel B is subtracted from panel A. Two regions in the spectrum were identified in which the empirical data had significantly more windows with significantly less (lower ellipse) and more (upper ellipse) Neanderthal ancestry than expected. Only windows with an expected introgression frequency greater than zero, less than 50% masked sites, intermediate recombination rate (i.e., ≥0.65 cm/Mb and ≤1.52 cM/Mb), and that have at least 50% African-like, at least 10% European-like, and less than 5% East Asian/Native American-like ancestry were included in these analyses. Densities and residuals were normalized to a range between 0 and 1. See also Figures S8 and S9.

**Fig. 5.**
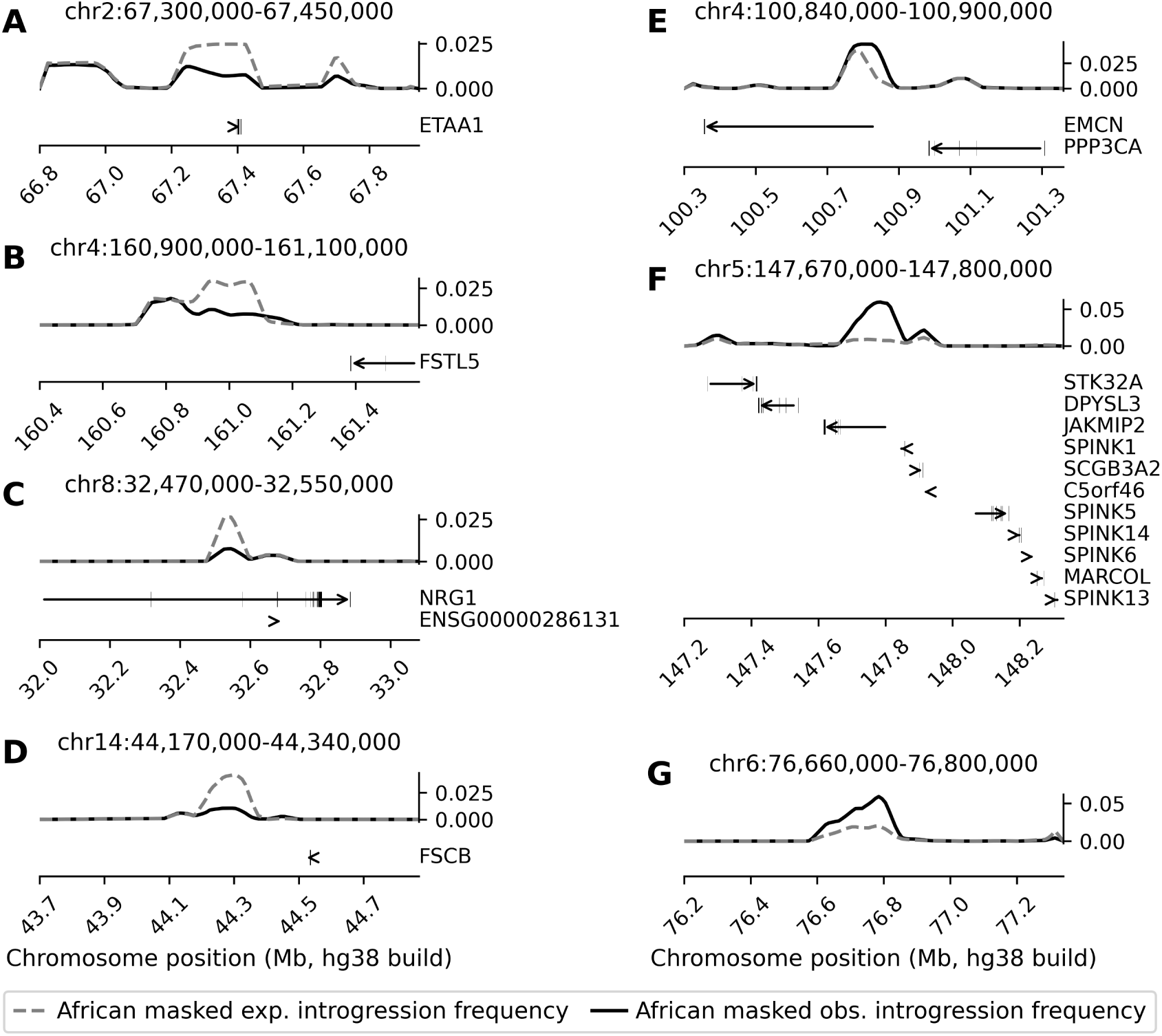
Expected and observed Neanderthal introgression frequencies as well as the localization of protein-coding genes within 500 kb in regions with significantly less (**A-D**) and more (**E-G**) Neanderthal ancestry than expected. Expected and observed introgression frequencies were calculated based on the African masked call set. Positions of the genomic regions with significantly less or more Neanderthal DNA than expected are shown in each panel. Gene locations were taken from GENCODE v46 (Frankish et al., 2023), and genomic positions are in hg38. See also Table S1.

The region with significantly less Neanderthal ancestry than expected on chromosome 2 overlaps with *ETAA1* (Figure 5A), which encodes a stress response protein that promotes DNA replication fork progression and integrity (Bass et al., 2016) and is active during mitosis and meiosis (Saldivar et al., 2018; Pereira et al., 2020). The depleted region on chromosome 4 is approximately 300 kb downstream of *FSTL5* (Figure 5B). The calcium ion-binding protein encoded by this gene is expressed in the brain (Lonsdale et al., 2013) and is associated with cancer (Remke et al., 2011; Zhang et al., 2015) but also obsessive-compulsive personality disorder (Lisboa et al., 2019). The depleted region on chromosome 8 overlaps with the *NRG1* (Figure 5C), encoding a glycoprotein that mediates cell-cell signaling, among others. *NRG1* is more ubiquitously expressed (Lonsdale et al., 2013) and has been implicated in schizophrenia (Stefansson et al., 2002). Furthermore, the depleted region on chromosome 14 is approximately 170 kb downstream of *FSCB* (Figure 5D). *FSCB* encodes a fibrous sheath CABYR-binding protein that is involved in spermatogenesis (Li et al., 2007). While the regions with significantly more Neanderthal ancestry than expected on chromosome 4 and chromosome 5 overlap with multiple genes (Figure 5E & F), the enriched region on chromosome 6 is not in the proximity of a protein-coding gene (Figure 5G). The enriched region on chromosome 4 overlaps with *ECMN* (also known as *MUC14*) and is approximately 125 kb downstream of *PPP3CA* (Figure 5E). *ECMN* inhibits cell adhesion and cell interactions with extracellular matrix (Kinoshita et al., 2001). The enriched region on chromosome 5 overlaps with *JAKMIP2*, which is part of the Golgi apparatus and expressed in brain tissues (Lonsdale et al., 2013), but it is also in proximity to several other genes (Figure 5F), including members of the *SPINK* gene family that are involved in innate immunity (Rimphanitchayakit and Tassanakajon, 2010).

We also compared expected and observed introgression frequencies in the 30,780 admixed individuals for 93 previously identified candidate loci of adaptive Neanderthal introgression in European populations (Racimo et al., 2017) (see Materials and Methods). These loci did not overlap with identified outlier regions in this study as they generally had introgression frequencies that matched expectations based on local ancestry patterns and introgression frequencies in the reference populations. However, three loci have a higher Neanderthal introgression frequency than would be expected after 15 generations of drift: chr5:168,652,996-168,692,995, chr9:16,800,003-16,840,002, and chr18:53,993,631-54,033,630. The region on chromosome 5 overlaps with *SLIT3*, and the region on chromosome 9 overlaps with *BNC2*. *SLIT3* is associated with tumor suppression (Marlow et al., 2008), while *BNC2* is the classical example of adaptive introgression and is associated with skin pigmentation, among others (Reilly et al., 2022).

### 2.4 Hybrid incompatibilities and intrinsic negative selection have shaped introgression landscapes

Previously, large introgression deserts have been described in Eurasian populations (Sankararaman et al., 2014; Vernot and Akey, 2014; Vernot et al., 2016; Sankararaman et al., 2016; Chen et al., 2020). However, the evolutionary mechanisms leading to these deserts are still debated with hybrid incompatibilities (Sankararaman et al., 2014, 2016; Harris et al., 2023) and intrinsic negative selection (Juric et al., 2016; Vernot et al., 2016; Harris and Nielsen, 2016; Kim et al., 2018; Steinrücken et al., 2018; Petr et al., 2019) as non-mutually exclusive explanations. With respect to hybrid incompatibilities being the cause, it has been hypothesized that genetic incompatibilities reduced hybrid fertility (Jégou et al., 2017). If there are novel desert-like regions in admixed individuals, their evolutionary genetics may allow disentangling of these hypotheses.

To identify novel introgression desert-like regions, we searched for large genomic regions (i.e., ≥8 Mb) that contain significantly less Neanderthal DNA than expected using the African masked call set of Neanderthal introgressed segments (Equation 3). We identified four emerging deserts on chromosomes 2, 7, 10, and 17. The novel desert-like region on chromosome 7 overlapped with a known Neanderthal introgression desert (Vernot et al., 2016; Chen et al., 2020), and for this reason, was excluded from subsequent analyses (Figure 6A; Table S2). We note that we did not observe any novel desert-like regions in simulations. To confirm that the three novel desert-like regions are under background selection, we evaluated B-statistics (McVicker et al., 2009). Indeed, we found that the three novel desert-like regions have lower B-statistics compared to the genome-wide background and previously known deserts (Mann-Whitney U *p* ≤ 10^−6^ and *p* ≤ 10^−6^, respectively; Figure 6B; Table S2), indicating stronger background selection.

**Fig. 6.**
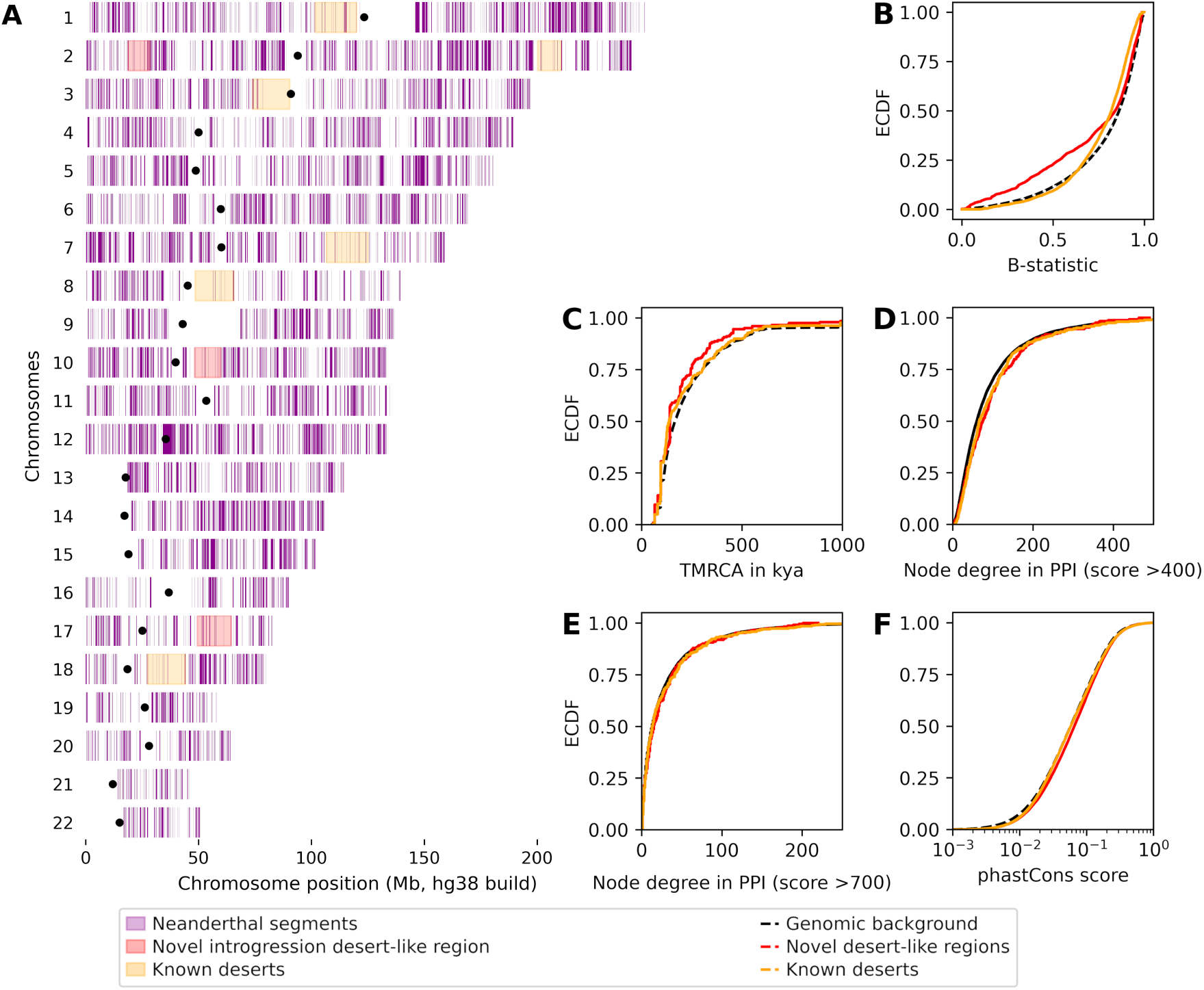
The localization and evolutionary genetics of novel introgression desert-like regions and previously known deserts. **A)** The genome-wide distribution of African masked Neanderthal haplotypes (purple) and the localization of novel desert-like regions (red) and previously known introgression deserts (orange) (Vernot et al., 2016; Chen et al., 2020). Genomic positions are in hg38. **B)** Novel introgression desert-like regions are subject to stronger background selection (lower B-statistic) than the genome-wide background and previously known deserts (Mann-Whitney U *p* ≤ 10^−6^ and *p* ≤ 10^−6^). Previously known deserts are also subject to stronger background selection than the genome-wide background (Mann-Whitney U *p* ≤ 10^−6^). **C)** Neanderthal-derived alleles in novel introgression desert-like regions and previously known introgression are younger than expected by chance (Mann-Whitney U *p* = 6.47 × 10^−4^ and *p* = 8.15 × 10^−6^). **D)** Genes overlapping the novel desert-like regions and previously known deserts interact with slightly more proteins than random genes (*p* = 2.05 × 10^−3^ and *p* = 1.37 × 10^−3^) when considering medium confidence protein-protein interactions in STRING (i.e., score *>* 400). **E)** The shifts for genes with more interactions disappear when only considering high-confidence interaction in STRING (score *>* 700; *p* = 0.28 and *p* = 0.42). **F)** Novel introgression desert-like regions and previously known deserts show a small shift towards greater phastCons scores compared to the genome-wide background (Mann-Whitney U *p* ≤ 10^−6^ and *p* ≤ 10^−6^). See also Tables S2 and S3.

To test whether the evolution of desert-like regions is driven by hybrid incompatibilities or intrinsic negative selection, we interrogated these novel desert-like regions and previously known deserts from Vernot et al. (2016) and Chen et al. (2020) (Table S2) for several evolutionary genetic statistics. First, we compared the allele ages of Neanderthal-derived variants in these regions, i.e., variants present in one or more Neanderthal reference genomes but absent from the Denisovan genomes and African reference genomes. Neanderthal-derived variants in novel desert-like regions and previously known deserts are modestly younger than the genomic background (Mann-Whitney U *p* = 6.47 × 10^−4^ and *p* = 8.15 × 10^−6^, respectively; Figure 6C; see Supplemental Information), making them more likely to be epistatically incompatible in a human genetic background (see Discussion). Furthermore, genes overlapping these novel desert-like regions and previously known deserts also interact with slightly more proteins than random genes when considering all interactions with at least medium confidence (Mann-Whitney U *p* = 2.05 × 10^−3^ and *p* = 1.37 × 10^−3^, respectively; Figure 6D) but do not have more interaction partners than random genes when only considering high-confidence interactions (Mann-Whitney U *p* = 0.28 and *p* = 0.42, respectively; Figure 6E). A gene set enrichment analysis also revealed that the three novel desert-like regions are nominally enriched for genes associated with reproductive processes (GO:0022414; FDR-controlled *p* = 0.052), among others (Table S3). However, we also observed a small but statistically significant shift towards larger phastCons scores (Siepel et al., 2005) in novel desert-like regions and known introgression deserts compared to the genome-wide background (Mann-Whitney U *p* ≤ 10^−6^ and *p* ≤ 10^−6^, respectively; Figure 6F), indicating greater evolutionary conservation and that intrinsic negative selection is more likely to remove Neanderthal DNA from these regions. Thus, both hybrid incompatibilities and intrinsic negative selection may have shaped introgression deserts in modern human genomes.

## 3 Discussion

Leveraging 30,780 admixed genomes with predominantly recent African-like and European-like ancestry, we found no evidence for strong, polygenic selection of Neanderthal introgressed segments that were brought into an African genetic background during the past 15 generations since admixture. When focusing on Neanderthal segments mostly contributed by European-like ancestors, admixed genomes contain approximately as much Neanderthal ancestry as expected based on continental ancestry proportions and average amounts of Neanderthal DNA in each of these source ancestries (Figure 3). This is consistent with previous studies showing that the amount of Neanderthal ancestry in modern human genomes has been constant for the past 45,000 years and that archaic haplotypes do not carry more deleterious variants than non-archaic haplotypes (Harris and Nielsen, 2016; Dannemann et al., 2017; Petr et al., 2019; Telis et al., 2020; Skov et al., 2020).

Yet, Neanderthal ancestry may still be under selection in local genomic regions. After accounting for drift by conditioning on the simulated joint spectrum of expected and observed introgression frequencies in 50 kb windows, we identified four and three independent genomic regions with significantly less and more Neanderthal ancestry than expected, respectively (Figure 5; Table S1). We note that by looking for less or more Neanderthal ancestry within recent European-like ancestry tracts our evolutionary analysis in admixed populations is complementary to previous work that examined local ancestry proportions to infer whether there was evidence of strong natural selection following the middle passage (Bhatia et al., 2014) and searched for signatures of adaptive introgression in Eurasian populations (Racimo et al., 2017; Gittelman et al., 2016). Previously identified candidate loci of adaptive introgression in European populations had introgression frequencies that matched expectations in admixed individuals (Racimo et al., 2017), suggesting that they have not been under strong positive selection during the last 15 generations. Furthermore, genetic features of significant outlier regions in our study are consistent with earlier findings that some of the strongest signals of adaptive introgression are in genes related to immunity (Reilly et al., 2022; Zeberg et al., 2024). For example, one of the identified regions with significantly more Neanderthal ancestry than expected in this study (chr5:147,670,000-147,800,000) is in the proximity of several members of the *SPINK* gene family that are associated with innate immunity (Figure 5F; Table S1). We point out that this region has a complex evolutionary history with a 2 million-year-old deletion in the nearby *STK32A* gene and a *>*1.5 million-year-old inversion in *SPINK14* that have recently been identified as candidates of selective pressures on the lineage leading to modern humans (Aqil et al., 2023; Giner-Delgado et al., 2019).

Another longstanding question of Neanderthal introgression is whether hybrid incompatibilities or intrinsic negative selection against Neanderthal ancestry led to the formation of large introgression deserts (Sánchez-Quinto and Lalueza-Fox, 2015; Reilly et al., 2022). To disentangle the hypotheses of hybrid incompatibilities and intrinsic negative selection, we compared evolutionary genetic statistics of three newly identified desert-like regions and previously known deserts (Figure 6; Table S2), including estimated ages of Neanderthal-derived variants. Hybrid incompatibilities can arise from multiple mutations on the same lineage, i.e., ancestral-derived incompatibilities (Wang et al., 2013). Due to the snowball effect (Orr, 1995), one would expect mutations on the Neanderthal branch that occurred long after the human-Neanderthal split, i.e., younger Neanderthal-derived alleles, to be more likely to result in ancestral-derived hybrid incompatibilities. Indeed, we found that Neanderthal-derived variants in introgression desert-like regions and known deserts are younger than in other parts of the genomes (Figure 6C), and their potential to be genetically incompatible is further compounded by the colocalization with connected genes in these regions (Figure 6D & E). Furthermore, we found that these desert-like regions are nominally enriched for genes involved in reproductive processes (Table S3). Given that we identified short regions with significantly less Neanderthal ancestry than expected in the proximity of genes involved in spermatogenesis (*FSCB*) and mitosis/meiosis (*ETAA1*), among others, the depletion of Neanderthal ancestry around reproductively important genes appears to be a general pattern. Such a depletion pattern fits with the hypothesis that genetic incompatibilities in reproductively relevant genes reduced hybrid fertility (Sankararaman et al., 2014, 2016; Jégou et al., 2017). However, hybrid incompatibilities are not mutually exclusive from intrinsic negative selection against Neanderthal ancestry in these regions. We also observed a higher evolutionary constraint in these regions (Figure 6D), which makes negative selection more likely to remove Neanderthal-derived variants. The desert-like region on chromosome 10 overlaps with *BICC1*, a gene that was previously identified as a candidate for positive selection in early modern humans (Green et al., 2010). This indicates that the evolutionary dynamics in these regions may be heterogeneous, and different evolutionary forces may have acted on them. Therefore, these regions require further study to fully understand their evolutionary histories.

Our study is not without limitations. Since admixture occurred only 15 generations ago, selection on Neanderthal haplotypes would have had to be strong for us to be able to detect it. We found that null expectations from our probabilistic modeling of expected Neanderthal introgression frequencies in 50 kb windows were not well calibrated to identify windows with significantly less or more Neanderthal ancestry than expected in the admixed genomes, despite efforts to account for 15 generations of drift (Equation 3 and Supplemental Information; Figure S9). This is possibly because our model does not capture effects from deeper population history, e.g., the out-of-Africa bottleneck. For this reason, we took a more conservative approach and conditioned our identification of short regions with significantly less and more Neanderthal ancestry than expected on the simulated joint spectrum of expected and observed introgression frequencies. Furthermore, we do not know the exact ancestry composition of the admixing populations 15 generations ago. However, as total amounts of Neanderthal ancestry (Figure 2A) and Neanderthal introgression frequencies per 50 kb windows (Figure S11) are very similar across populations from the same continental ancestry group, it is unlikely that this is a significant confounder. Nevertheless, identified regions that are putatively under selection require further validation.

In summary, we showed that the remaining Neanderthal ancestry appears to be largely evolutionary neutral in contemporary genomes, that is, we did not find evidence for strong, polygenic selection of Neanderthal ancestry in admixed genomes with African-like ancestry. Furthermore, we uncovered additional evidence for the potential involvement of hybrid incompatibilities in shaping the introgression landscapes of our species.

## 4 Materials and Methods

### 4.1 Materials availability

This study did not generate new unique reagents.

### 4.2 Data and code availability

This study used data from the All of Us Research Program’s Controlled Tier Dataset v7.1, which is available to authorized users on the Researcher Workbench and publicly available data from the 1000 genomes project phase 3. All analyses described above have been implemented in a Snake-make workflow (Mölder et al., 2021). All code used and computed introgression and local ancestry frequencies are available from https://github.com/LachanceLab/introgression in admixed genomes.

### 4.3 Method Details

#### 4.3.1 Dataset description

##### Ethics statement

All study participants in the All of Us Research Program provided written consent in accordance with the Declaration of Helsinki and the U.S. Common Rule. As per Georgia Institute of Technology IRB protocol H15385, all genomic data analyzed in this study was deidentified. The authors declare no conflicts of interest.

##### Modern human samples

Using previously estimated ancestry proportions (Conley et al., 2023; All of Us Research Program Investigators et al., 2019; Bick et al., 2024), we identified 30,780 unrelated recently admixed individuals who had at least 50% African-like ancestry, at least 10% European-like ancestry, and at most 5% East Asian/Native American-like ancestry and for whom short-read whole-genome sequences are available in All of Us v7.1 (All of Us Research Program Investigators et al., 2019; Bick et al., 2024). As we considered continental ancestry proportions, we aggregated inferred East Asian-like and Native American-like ancestry proportions. By limiting the analyses to mostly two-way admixed individuals, we aimed to improve the interpretability of the empirical dynamics. For computational reasons, we then used the ACAF v7.1 genotype call set that only includes variants that have a population-specific allele frequency ≥100 or a population-specific allele count ≥100 in any All of Us computed ancestry group. This call set contains 48,314,438 variable sites and 99,250,816 variants. For all analyses described below, we only considered autosomal data.

We constructed continental reference panels of introgression landscapes using 1000 genomes project (1KGP) phase 3 (Auton et al., 2015) and All of Us v7.1 (All of Us Research Program Investigators et al., 2019; Bick et al., 2024). Specifically, we used 1KGP populations that are assigned to African (504 individuals, i.e., excluding admixed ACB & ASW), European (503 individuals), and East Asian (504 individuals) superpopulations. 1KGP populations assigned to the East Asian super-population were used as a proxy to characterize the introgression landscape in Native American genomes, which were previously shown to have similar levels of Neanderthal introgressed sequence per individual (Sankararaman et al., 2016). 1KGP genotype calls were lifted over from hg19 to hg38 coordinates using CrossMap v0.6.5 (Zhao et al., 2013). To obtain more granular estimates of introgression frequency, we added 563 unrelated individuals with ≥ 99% African-like ancestry (AOU-AFR), 10,000 random, unrelated (i.e., no first- or second-degree relatives) individuals with ≥ 99% European-like ancestry (AOU-EUR), and 71 unrelated individuals with ≥ 99% Native American-like ancestry (AOU-NA) from All of Us using previously estimated continental ancestry proportions (Conley et al., 2023; All of Us Research Program Investigators et al., 2019; Bick et al., 2024). In sum, the reference panels included 1,067 individuals with African-like ancestry, 10,503 individuals with European-like ancestry, and 575 individuals with East Asian/Native American-like ancestry.

##### Archaic hominin reference genomes

We used all three high-quality Neanderthal reference genomes available to date, i.e., the Altai, Vindija33.19, and Chagyrskaya individual (Prüfer et al., 2013, 2017; Mafessoni et al., 2020), as well as the Denisovan reference genome (Meyer et al., 2012). In the main text, we focus on results using the Vindija33.19 individual because its genome is the closest to the introgressing Neanderthal lineage (Prüfer et al., 2017; Mafessoni et al., 2020). However, we note that all available Neanderthal reference genomes yield qualitatively similar results on a smaller test set of 93 admixed individuals from 1KGP-ACB & 1KGP-ASW (see Supplemental Information and Figure S7). All genotype calls and filters were lifted over from hg19 to hg38 human reference genome using CrossMap v0.6.5 (Zhao et al., 2013).

#### 4.3.2 Detection of Neanderthal introgressed tracts

We chose IBDmix v1.0.1 to detect introgressed segments (Chen et al., 2020). Note that IBDmix does not require an unadmixed reference panel. We followed the procedure described in the original publication but applied a more stringent mask. Specifically, we applied the following filters when calling introgressed segments:

- Recommended minimal filter mask for the respective archaic genome (Meyer et al., 2012; Prüfer et al., 2013, 2017; Mafessoni et al., 2020). The masks were downloaded from http://cdna.eva.mpg.de/neandertal/.
- We determined mappable regions, i.e., the majority of 35-mers are mapped uniquely without 1-mismatch to the hg38 reference genome (i.e., Heng Li’s SNPable regions mask) (Li and Durbin, 2011).
- We excluded regions that were predicted to be introgressed from modern humans into Neanderthals with 90% probability by ArgWeaver-D (Hubisz et al., 2020).
- We removed segmental duplications, repetitive regions, and gaps in the hg38 assembly. These files were downloaded from the USCS Table Browser (Karolchik et al., 2004).
- We excluded sites inaccessible in 1KGP data and sites within 5 bp of indels in 1KGP data (Auton et al., 2015).
- We removed CpG sites as per Vernot and Akey (2014) using African 1KGP reference population as well as chimpanzee (panTro6), bonobo (ponAbe3), and rhesus macaque (rheMac10) reference genomes.

Applying the above mask, introgressed segments were then separately called for each population, i.e., reference subpopulations and admixed individuals, to avoid confounding from population structure. Following Chen et al. (2020), we only retained introgressed segments that were at least 50 kb long and had a LOD score of at least 4.0. To account for ILS, we then refined Neanderthal call sets by filtering out segments that overlapped with a Denisovan introgressed segment in an African reference individual by at least 1 bp using bedtools v2.30.0 (Quinlan and Hall, 2010). We refer to this call set as the “unfiltered” call set.

To account for remaining biases from ILS and false positive predictions, we filtered out additional Neanderthal introgressed segments. First, we additionally removed any introgressed segments that overlapped with a predicted segment in an African reference genome by at least 1 bp, i.e., an “African mask”. The reasoning for this is twofold: i) despite including ArgWeaver-D predicted human-to-Neanderthal introgressed regions in the mask (Hubisz et al., 2020) and using the Denisovan genome to control for ILS, IBDmix still has a higher false-positive rate in an African genetic background due to earlier human-to-Neanderthal introgression events (Harris et al., 2023; Li et al., 2024), and ii) regardless of whether a predicted introgressed segment in an African genome is a true or false positive prediction, we are most interested in segments not previously found in African genomes. This is because only the evolutionary fate of those segments has been assessed in an African genetic background in the admixed genomes during the last 15 generations. All analyses are based on this African masked call set of Neanderthal introgressed segments. Furthermore, to account for potential effects from recombination rate variation along the genome, we applied a recombination mask (Figure S5). That is, following Harris et al. (2023), we calculated the average recombination rate in non-overlapping 300 kb windows using the hg38 HapMap recombination map (Frazer et al., 2007). We then only retained windows with intermediate recombination rates, i.e., a recombination rate ≥0.65 cm/Mb (33*^rd^* percentile) and ≤ 1.52 cm/Mb (66*^th^* percentile). Finally, we only retained segments/50 kb windows that were fully covered by retained 300 kb windows using bedtools v2.30.0 (Quinlan and Hall, 2010).

#### 4.3.3 Local ancestry inference

We first phased genotype calls for the 30,780 recently admixed individuals using Beagle v5.4 with default parameters (Browning et al., 2021) and subsequently inferred local ancestry using FLARE v0.5.1 (Browning et al., 2023). As recommended by the authors, FLARE was trained on chromosome 1 using the above-mentioned 1KGP continental reference populations, i.e., 1KGP-AFR, 1KGP-EUR, and 1KGP-EAS superpopulations, and the trained model was subsequently used to infer local ancestries on the remaining autosomes. For each chromosome, we used the respective HapMap hg38 recombination map (Frazer et al., 2007). Figure S12 shows that genome-wide ancestry estimates by FLARE are highly concordant with the previously inferred ancestry proportions (Conley et al., 2023; All of Us Research Program Investigators et al., 2019; Bick et al., 2024); with Pearson’s correlations of *r* = 0.99 for recent African-like and European-like ancestry estimates. For both phases, local ancestry tracts were then extracted from the obtained output VCF files. Given a set of consecutive variants with the same assigned local ancestry, we defined a local ancestry track as the genomic region delimited by the positions of the first and last such variant.

#### 4.3.4 Genome-wide modeling of expected amounts of Neanderthal introgressed sequence per individual

All Neanderthal introgressed sequences found in the 30,780 admixed individuals must have passed through one of the admixing populations. Therefore, the introgression landscape in the admixed individuals is a function of recent ancestry patterns and introgression landscapes in the admixing populations, assuming random inheritance and neutrality. Due to the lack of a sufficiently sized Native American reference panel, we modeled the contribution of Neanderthal ancestry from the Native American-like component to the admixed individuals using the summed inferred East Asian-like and Native American-like ancestry proportions and the combined introgression landscape in East Asian and Native African reference genomes from 1KGP and All of Us, respectively. For completeness, we always describe the models for three-way admixed individuals (i.e., an African-like, European-like, and East Asian/Native American-like component), but for analyses of the African masked call set of Neanderthal introgressed segments, the term for the African-like component is omitted.

Under neutrality, the expected amount of Neanderthal introgressed sequence per individual inherited from admixing population *j* is proportional to the admixture proportion (*Q_j_*) times the average amount of Neanderthal introgressed sequence per individual (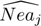) in the respective admixing population. Thus, for the admixed individuals with African-like (AFR), European-like (EUR), and East Asian/Native American-like (EAS/NA) ancestry, the expected amount of Neanderthal introgressed sequence per individual is given by:

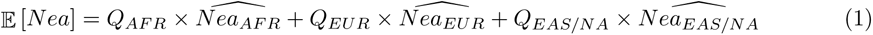

To test for a general depletion/enrichment of Neanderthal ancestry in admixed genomes, we fitted a linear least-square regression to the expected and observed amounts using the implementation in scipy v1.10.1 (Virtanen et al., 2020). Under neutrality, one would expect a slope of one and a y-intercept of zero.

#### 4.3.5 Modeling of expected Neanderthal introgression frequencies in 50 kb windows

To calculate expected introgression frequencies in the admixed individuals, we model the expected number of Neanderthal introgression haplotypes in the admixed population as binomial draws of Neanderthal haplotypes from the source populations. First, we segmented the genome into overlapping 50 kb windows (step size 10 kb) and computed local ancestry frequencies, i.e., frequencies of African-, European-, and East Asian/Native American-like haplotypes, in each window by calculating the average fraction of base pairs per window that are covered by tracts with a given ancestry across all admixed individuals and both phases, using bedtools v2.30.0 (Quinlan and Hall, 2010). Similarly, we calculated introgression frequencies for each source population in each window using the African masked call set of Neanderthal introgressed segments (*q_i,j_*). Second, to allow binomial sampling, we converted the frequency of tracts with recent ancestry *j* in window *i* (*n_i,j_*) to discrete numbers by multiplying them with the number of admixed individuals (*N_Adm_*). We imposed the constraint that the total number of local ancestry tracts must sum up to the number of admixed individuals, i.e., ∑*_j_*_∈{AFR, EUR, EAS/NA}_ [*n_i,j_N_Adm_*] = *N_Adm_* because we deal with pseudo-haploid genomes since IBDmix does not provide phase information. Third, to account for sampling error in observed Neanderthal introgression frequencies, we calculated binomial proportion confidence intervals according to Agresti-Coull (i.e., 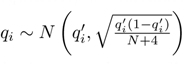, where 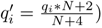) (Agresti and Coull, 1998). Finally, assuming random inheritance and neutrality and by integrating over the 99% Agresti-Coull binomial proportion confidence intervals of introgression frequencies for each ancestry component, the expected number of introgressed haplotypes overlapping window *i* in the admixed individuals (*X_i,Adm_*) is given by the following multinomial distribution:

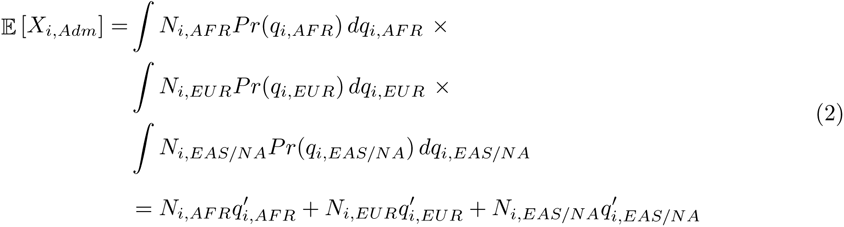

where *N_i,j_* is the number of haplotypes of recent ancestry *j* (i.e., *n_i,j_* × *N_Adm_*), *q_i,j_* is the estimated introgression frequency in admixing population *j*, and 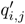 is the center-point adjusted Agresti-Coull estimate of the introgression frequency in admixing population *j* in window *i*.

#### 4.3.6 Probabilistic identification of 50 kb windows with significantly less and more Neanderthal ancestry than expected

The above-described model for the expected number of introgressed haplotypes (Equation 2) also allows calculating probabilities of observed frequencies being significantly lower or higher than expected in the admixed genomes. We first calculated the 95% Agresti-Coull binomial proportion confidence interval of introgression frequencies in the admixed genomes for each window *i* as described above. We then converted this confidence interval of introgression frequencies to a range of discrete numbers of Neanderthal introgressed haplotypes in the admixed individuals by multiplying them with the number of admixed individuals and taking the floor (i.e., *X_i,Adm_* = ⌊*q_i,Adm_N_Adm_*⌋). Note that taking the ceiling yields qualitatively similar results. For windows with a lower Neanderthal introgression frequency than expected, we calculated the probability of observing a lower Neanderthal introgression frequency than given by the upper bound of the 95% Agresti-Coull interval 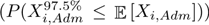. In contrast, for windows with a higher Neanderthal introgression frequency than expected, we calculated the probability of observing a higher Neanderthal introgression frequency than given by the lower bound of the 95% Agresti-Coull interval 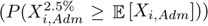. For example, the probability of observing a lower Neanderthal introgression frequency than expected is given by:

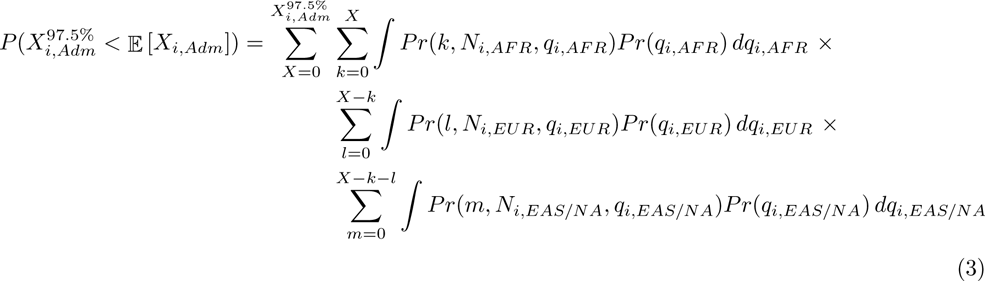

where the outer sum accounts for contributions less than or equal to 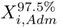, the inner sums account for all possible combinations of Neanderthal haplotype contributions from the different admixing populations that add up to *X*, and the integrals account for uncertainties in the estimated introgression frequency in the respective reference populations. Similarly, 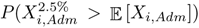 is calculated for windows that show higher introgression frequencies than expected by setting the limits of the outer sum to 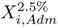 and *N_Adm_*, respectively. That is, given the data, we calculated the probability of the weakest plausible depletion or enrichment, respectively.

We note that we found Equation 3 identifies numerous false positive 50 kb windows with significantly less or more Neanderthal ancestry than expected based on local ancestry patterns and introgression frequencies in the reference populations despite attempts to account for 15 generations of drift (see Supplemental Information; Figure S9). However, we found that Equation 3 is well calibrated for identifying large genomic regions (≥ 8 Mb) with significantly less Neanderthal ancestry than expected (see below).

#### 4.3.7 Identifying genomic 50 kb windows with significantly less or more Neanderthal ancestry than expected under neutral evolution

We found that probabilistic modeling under the approved described model of binomial inheritance was not well calibrated to identify windows with significantly less or more Neanderthal ancestry than expected as numerous significant windows were identified in neutral simulations (see Equation 3 and Supplemental Information; Figure S9). For this reason, we took an alternative, more conservative approach and controlled for genetic drift by conditioning on the simulated joint spectrum of expected and observed introgression frequencies (see below for details on the simulations). We calculated the joint spectrum of expected and observed introgression frequency based on windows with expected introgression frequency greater than zero, less than 50% masked sites, intermediate recombination rate (i.e., 0.65 cm/Mb ≤ recombination rate ≤ 1.52 cm/Mb; see above), and that have at least 50% African-like, at least 10% European-like, and less than 5% East Asian/Native American-like ancestry. We binned the windows into 0.002 frequency bins, and calculated the fraction of windows falling into a given bin for the empirical data and aggregated the data from all simulation replicates. We then subtracted the simulated joint spectrum from the empirical joint spectrum and applied a Gaussian smoothing filter (*σ* = 2 and *radius* = 6 bins) to the resulting residual spectrum. Subsequently, we calculated the Euclidean distance of the smoothed residuals to the background level and identified local maxima with a minimum intensity of 10^−5^. The identified local maxima were used to seed the Watershed algorithm for detecting peaks in the residual spectrum. Finally, we only considered windows with significantly less or more Neanderthal ancestry than expected falling into identified peak regions and merged windows within 50 kb from each other using bedtools v2.30.0 (Quinlan and Hall, 2010). All peak detection steps were implemented using scikit-image v0.23.2 (Walt et al., 2014).

#### 4.3.8 Characterizing previously identified candidate loci of adaptive introgression in the admixed population

A previous scan for adaptive Neanderthal introgression by Racimo et al. (2017) identified several candidate loci for adaptive introgression (Table S3 in Racimo et al. (2017)). We selected all loci that were identified as candidate loci for adaptive introgression from Neanderthals or Neanderthals and Denisovans in individual European populations, a European continental target panel, or a Eurasian target panel, yielding 370 candidate loci. The coordinates of these candidate loci were lifted over from hg19 to hg38 coordinate system using CrossMap v0.6.5 (Zhao et al., 2013). These regions were then intersected with 50 kb windows that had evidence of Neanderthal introgression in the admixed individuals after applying the African mask, i.e., after removing Neanderthal introgressed segments overlapping with an introgressed segment in African reference genomes, using bedtools v2.30.0 (Quinlan and Hall, 2010). 93 out of 370 candidate loci overlapped with 50 kb windows had introgression frequencies greater than zero in the admixed using the African masked call set of Neanderthal introgressed segments. Finally, we intersect these 93 regions with identified outlier regions in this study and compared expected and observed introgression frequencies in the admixed individuals.

#### 4.3.9 Identifying novel Neanderthal introgression desert-like regions

Previous studies characterizing the Neanderthal introgression landscape in modern Eurasian genomes identified large genomic regions (≥ 8 Mb) that are significantly depleted for Neanderthal DNA, so-called introgression deserts (Sankararaman et al., 2014; Skov et al., 2020; Chen et al., 2020). Taking a similar approach as Chen et al. (2020), we searched for large genomic regions with significantly less Neanderthal ancestry than expected in the admixed genomes by segmenting the genome into overlapping windows of various sizes (8 − 15 Mb), using a step size of 100 kb. For each window, we first quantified the amount of introgression by summing the number of base pairs of overlapping introgressed segments across all individuals and normalizing by the window size, excluding windows in which ≥ 50% of the sites were masked. We then only searched for emerging introgression desert-like regions to windows with introgression frequencies in the bottom 5*^th^* percentile in admixed genomes for each window size and identified windows that are significantly depleted for Neanderthal ancestry in the admixed population relative to the reference populations at a Bonferroni corrected significance level of 0.05, using Equation 3. Equation 3 is well calibrated for this purpose since we did not identify any large genomic regions with significantly less Neanderthal ancestry than expected in neutral simulations. Finally, overlapping windows were merged using bedtools v2.30.0.

To disentangle different hypotheses for the evolutionary origin of these deserts, we annotated newly emerging introgression desert-like regions with various evolutionary statistics. Identified introgression desert-like regions and previously known introgression desert in Eurasian genomes (Vernot et al., 2016; Chen et al., 2020) were annotated with B-statistics (McVicker et al., 2009), phastCons scores (Siepel et al., 2005), estimated allele ages of non-CpG Neanderthal-derived variants (see Supplemental Information), and number of protein-protein interactions in the STRING database v11.5 of overlapping genes (Szklarczyk et al., 2019). We analyzed the number of interaction partners in STRING using two confidence score cutoffs for interactions: i) medium confidence (score *>*400) and ii) high confidence (score *>*700). With respect to these summary statistics novel desert-like regions and previously known deserts were then compared to the genomic background, and statistical significance was assessed using a Mann-Whitney U test, as implemented in scipy v1.10.1 (Virtanen et al., 2020). As B-statistics and phastCons scores are calculated for short intervals, we weighted them by overlap with the regions of interest. Furthermore, we conducted a gene set enrichment analysis of genes overlapping novel introgression desert-like regions using DAVID (Sherman et al., 2022). The Functional Annotation Tool was used to identify enriched GO terms, and the false discovery rate was controlled using Benjamini-Hochberg.

#### 4.3.10 Neutral simulations of ancient introgression and recent admixture

We performed neutral coalescence simulations, using msprime v1.2.0 (Baumdicker et al., 2021), to ensure that our above-described approaches for testing for secondary selection of Neanderthal alleles are well calibrated.

We extended the three populations out-of-African (OOA) model by Gravel et al. (2011) to include archaic introgression and recent admixture in the Americas (Figure S13). Specifically, assuming a generation time of 25 years, we simulated an ancestral population with an effective population size (*N_e_*) of 7,310, from which a Neanderthal population split off 28,000 generations ago. Subsequently, a Denisovan population split off from the Neanderthal population 20,000 generations ago. *N_e_*for the Neanderthal was set to 2,800 and *N_e_* for the Denisovan populations was set to 2,600. With the emergence of anatomically modern humans 5,920 generations ago, the *N_e_* of the African population was expanded to 14,474. We simulated the OOA migration 2,040 generations ago, and the OOA population experienced a bottleneck with a *N_e_* of 1,861. We simulated symmetric migration between the African population and the OOA population at a rate of 1.5 × 10^−4^ per generation. The OOA population then received a 5% Neanderthal introgression pulse 1,500 generations ago, prior to the split of the European and East Asian populations. This split was simulated to have occurred 920 generations ago. The European and East Asian populations experienced an additional bottleneck with a *N_e_* of 1,032 for the European population and a *N_e_* of 554 for the East Asian populations, but then they grew exponentially with rates of 3.8×10^−3^ and 4.8×10^−3^ per generation, respectively. We simulated symmetric migration between the African and the European and East populations at rates of 2.5×10^−5^ and 7.8×10^−6^ per generation, respectively. Symmetric migration between the European and the East Asian populations was simulated at a rate of 3.11 × 10^−5^ per generation. Lastly, we simulated American admixture 15 generations ago with the following admixture proportions for African, European, and East Asian-like ancestry: 0.8, 0.19, and 0.1, respectively. Following Browning et al. (2018), the initial *N_e_* of the admixed population was set to 30,000, and post admixture, the admixed population was simulated to grow with a rate of 0.05 per generation (see Figure S13).

We simulated genomes with ten chromosomes each of which used hg38 HapMap recombination map for chromosome 16 (Frazer et al., 2007) and a mutation rate of 2.36 × 10^−8^ per base pair per generation. We performed ten replicate simulations, sampling the same number of reference and admixed individuals as considered in the empirical analyses (i.e., 1,067 African, 10,503 European, 575 Native American, and 30,780 admixed individuals). We used the discrete-time Wright-Fisher model (i.e., *dtwf*) in msprime to obtain realistic long-range genetic correlation and reduce the bias from the coalescent when sampling a large number of individuals (Nelson et al., 2020; Baumdicker et al., 2021).

The simulated data was written to VCF files. To estimate genome-wide ancestry proportions, we first computed the top 20 principal components based on LD-pruned biallelic SNPs with a minor allele frequency ≥0.01 for 504 African, 503 European, and 504 East Asian/Native American individuals using plink and plink2 (Chang et al., 2015). We then projected the admixed samples onto the reference principal component space and inferred global ancestry proportions using Rye v0.1 (Conley et al., 2023). When calling introgressed segments with IBDmix, we masked the low recombination rate region around the centromere (positions: 31,000,000 - 47,000,000) as this region led to the prediction of unreasonable long introgressed haplotypes (*>* 10 Mb). In all other aspects, the simulated data was analyzed using the same workflow as for the empirical data.

### 4.4 Key Resources Table

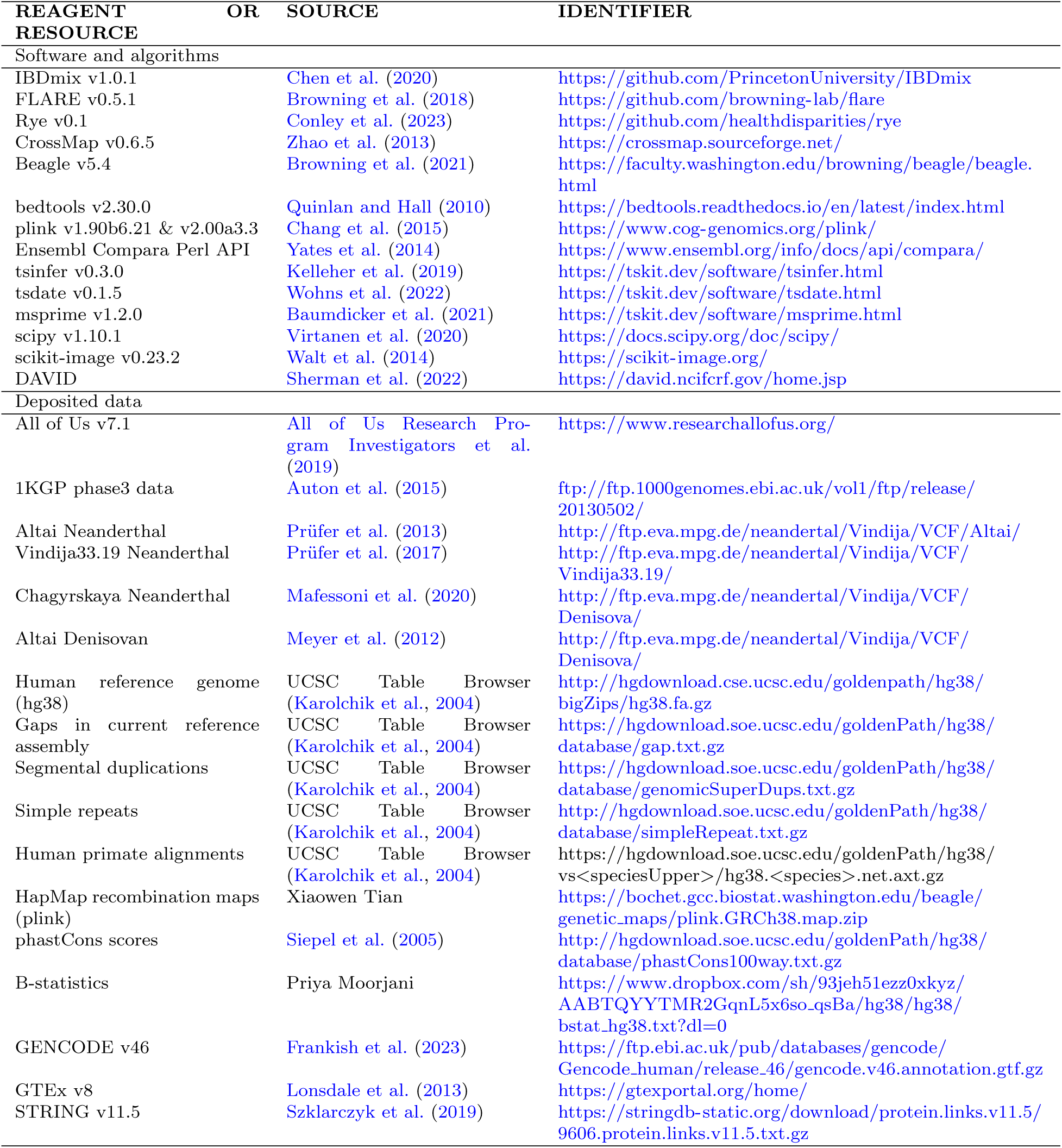

## Acknowledgments

We gratefully acknowledge All of Us participants for their contributions, without whom this research would not have been possible. We also thank the National Institutes of Health’s All of Us Research Program for making available the participant data examined in this study. We thank Shivam Sharma and I. King Jordan for sharing genetic ancestry estimates for All of Us participants. We thank Melissa Hubisz and Adam Siepel for providing files containing regions that were predicted to be introgressed from modern humans into archaic hominins for each of the used archaic reference genomes. We thank Priya Moorjani for providing files with pre-computed B-statistics to our lab. We thank Norman Johnson for valuable feedback on an earlier version of this manuscript. This research was supported in part through research cyberinfrastructure resources and services provided by the Partnership for an Advanced Computing Environment (PACE) at the Georgia Institute of Technology. This work was supported by the Google Cloud research credits program (GCP297878755) and an NIGMS MIRA grant to Joseph Lachance (R35GM133727).

## 5 Author Contributions

Conceptualization: A.P. and J.L.; Methodology: A.P. and J.L.; Formal Analysis: A.P.; Writing - Original Draft: A.P. and J.L.; Visualization: A.P. and J.L.; Supervision: J.L.; Funding Acquisition: J.L.

## Supplemental information

Document S1. Figures S1–S13 and Tables S1 - S3

## References

Agresti A, Coull BA (1998) Approximate is Better than “Exact” for Interval Estimation of Binomial Proportions. The American Statistician 52(2):119–126. 10.1080/00031305.1998.10480550

Aqil A, Speidel L, Pavlidis P, et al. (2023) Balancing selection on genomic deletion polymorphisms in humans. eLife 12:e79111. 10.7554/eLife.79111

Auton A, Abecasis GR, Altshuler DM, et al. (2015) A global reference for human genetic variation. Nature 526(7571):68–74. 10.1038/nature15393

Bass TE, Luzwick JW, Kavanaugh G, et al. (2016) ETAA1 acts at stalled replication forks to maintain genome integrity. Nature Cell Biology 18(11):1185–1195. 10.1038/ ncb3415

Baumdicker F, Bisschop G, Goldstein D, et al. (2021) Efficient ancestry and mutation simulation with msprime 1.0. Genetics 220(3). 10.1093/genetics/iyab229

Bhatia G, Tandon A, Patterson N, et al. (2014) Genome-wide Scan of 29,141 African Americans Finds No Evidence of Directional Selection since Admixture. The American Journal of Human Genetics 95(4):437–444. 10.1016/j.ajhg.2014.08.011

Bick AG, Metcalf GA, Mayo KR, et al. (2024) Genomic data in the All of Us Research Program. Nature pp 1–7. 10.1038/s41586-023-06957-x

Browning BL, Tian X, Zhou Y, et al. (2021) Fast two-stage phasing of large-scale sequence data. The American Journal of Human Genetics 108(10):1880–1890. 10.1016/j.ajhg.2021.08.005

Browning SR, Browning BL, Daviglus ML, et al. (2018) Ancestry-specific recent effective population size in the Americas. PLOS Genetics 14(5):1–22. 10.1371/journal.pgen.1007385

Browning SR, Waples RK, Browning BL (2023) Fast, accurate local ancestry inference with FLARE. The American Journal of Human Genetics 110(2):326–335. 10.1016/j.ajhg.2022.12.010

Chang CC, Chow CC, Tellier LCAM, et al. (2015) Second-generation PLINK: rising to the challenge of larger and richer datasets. GigaScience 4(1). 10.1186/s13742-015-0047-8

Chen L, Wolf AB, Fu W, et al. (2020) Identifying and Interpreting Apparent Neanderthal Ancestry in African Individuals. Cell 180(4):677–687.e16. 10.1016/j.cell.2020.01.012

Conley AB, Rishishwar L, Ahmad M, et al. (2023) Rye: genetic ancestry inference at biobank scale. Nucleic Acids Research 10.1093/nar/gkad149

Dannemann M, Prüfer K, Kelso J (2017) Functional implications of Neandertal introgression in modern humans. Genome Biology 18(1):1–11. 10.1186/s13059-017-1181-7

Frankish A, Carbonell-Sala S, Diekhans M, et al. (2023) GENCODE: reference annotation for the human and mouse genomes in 2023. Nucleic Acids Research 51(D1):D942–D949. 10.1093/nar/gkac1071

Frazer KA, Ballinger DG, Cox DR, et al. (2007) A second generation human haplotype map of over 3.1 million SNPs. Nature 449(7164):851–861. 10.1038/nature06258

Giner-Delgado C, Villatoro S, Lerga-Jaso J, et al. (2019) Evolutionary and functional impact of common polymorphic inversions in the human genome. Nature Communications 10(1):4222. 10.1038/s41467-019-12173-x

Gittelman RM, Schraiber JG, Vernot B, et al. (2016) Archaic Hominin Admixture Facilitated Adaptation to Out-of-Africa Environments. Current Biology 26(24):3375–3382. 10.1016/j.cub.2016.10.041

Gravel S, Henn BM, Gutenkunst RN, et al. (2011) Demographic history and rare allele sharing among human populations. Proceedings of the National Academy of Sciences of the United States of America 108(29):11983–11988. 10.1073/pnas.1019276108

Green RE, Krause J, Briggs AW, et al. (2010) A draft sequence of the neandertal genome. Science 328(5979):710–722. 10.1126/science.1188021

Harris DN, Platt A, Hansen ME, et al. (2023) Diverse african genomes reveal selection on ancient modern human introgressions in neanderthals. Current Biology 33(22):4905–4916.e5. 10.1016/j.cub.2023.09.066

Harris K, Nielsen R (2016) The Genetic Cost of Neanderthal Introgression. Genetics 203(2):881–891. 10.1534/genetics.116.186890

Hubisz MJ, Williams AL, Siepel A (2020) Mapping gene flow between ancient hominins through demography-aware inference of the ancestral recombination graph. PLOS Genetics 16(8):1–24. 10.1371/journal.pgen.1008895

Iasi LNM, Chintalapati M, Skov L, et al. (2024) Neandertal ancestry through time: Insights from genomes of ancient and present-day humans

Juric I, Aeschbacher S, Coop G (2016) The Strength of Selection against Neanderthal Introgression. PLOS Genetics 12(11):e1006340. 10.1371/journal.pgen.1006340

Jégou B, Sankararaman S, Rolland A, et al. (2017) Meiotic Genes Are Enriched in Regions of Reduced Archaic Ancestry. Molecular Biology and Evolution 34(8):1974–1980. 10.1093/molbev/msx141

Karolchik D, Hinrichs AS, Furey TS, et al. (2004) The ucsc table browser data retrieval tool. Nucleic Acids Research 32:D493–D496. 10.1093/nar/gkh103

Kelleher J, Wong Y, Wohns AW, et al. (2019) Inferring whole-genome histories in large population datasets. Nature Genetics 51(9):1330–1338. 10.1038/s41588-019-0483-y

Kim BY, Huber CD, Lohmueller KE (2018) Deleterious variation shapes the genomic landscape of introgression. PLOS Genetics 14(10):e1007741. 10.1371/JOURNAL.PGEN.1007741

Kinoshita M, Nakamura T, Ihara M, et al. (2001) Identification of human endomucin-1 and -2 as membrane-bound O-sialoglycoproteins with anti-adhesive activity. FEBS letters 499(1-2):121–126. 10.1016/s0014-5793(01)02520-0

Li H, Durbin R (2011) Inference of human population history from individual whole-genome sequences. Nature 475(7357):493–496. 10.1038/nature10231

Li L, Comi TJ, Bierman RF, et al. (2024) Recurrent gene flow between neanderthals and modern humans over the past 200,000 years. Science 385(6705):eadi1768. 10.1126/science.adi1768

Li YF, He W, Jha KN, et al. (2007) FSCB, a Novel Protein Kinase A-phosphorylated Calcium-binding Protein, Is a CABYR-binding Partner Involved in Late Steps of Fibrous Sheath Biogenesis. Journal of Biological Chemistry 282(47):34104–34119. 10.1074/jbc.m702238200

Lisboa BCG, Oliveira KC, Tahira AC, et al. (2019) Initial findings of striatum tripartite model in OCD brain samples based on transcriptome analysis. Scientific Reports 9(1):3086. 10.1038/s41598-019-38965-1

Lonsdale J, Thomas J, Salvatore M, et al. (2013) The Genotype-Tissue Expression (GTEx) project. Nature Genetics 45(6):580–585. 10.1038/ng.2653

Mafessoni F, Grote S, de Filippo C, et al. (2020) A high-coverage neandertal genome from chagyrskaya cave. Proceedings of the National Academy of Sciences 117(26):15132–15136. 10.1073/pnas.2004944117

Marlow R, Strickland P, Lee JS, et al. (2008) SLITs Suppress Tumor Growth In vivo by Silencing Sdf1/Cxcr4 within Breast Epithelium. Cancer Research 68(19):7819–7827. 10.1158/0008-5472.CAN-08-1357

McVicker G, Gordon D, Davis C, et al. (2009) Widespread Genomic Signatures of Natural Selection in Hominid Evolution. PLoS Genetics 5(5):1000471. 10.1371/journal.pgen.1000471

Meyer M, Kircher M, Gansauge MT, et al. (2012) A high-coverage genome sequence from an archaic denisovan individual. Science 338(6104):222–226. 10.1126/science.1224344

Mölder F, Jablonski KP, Letcher B, et al. (2021) Sustainable data analysis with Snakemake. Tech. Rep. 10:33, F1000Research, 10.12688/f1000research.29032.1

Nelson D, Kelleher J, Ragsdale AP, et al. (2020) Accounting for long-range correlations in genome-wide simulations of large cohorts. PLoS genetics 16(5):e1008619

Orr HA (1995) The Population Genetics of Speciation: The Evolution of Hybrid Incompatibilities. Genetics 139:180–185. URL https://academic.oup.com/genetics/article/139/4/1805/6013266

Pereira C, Smolka MB, Weiss RS, et al. (2020) ATR Signaling in Mammalian Meiosis: From Upstream Scaffolds to Downstream Signaling. Environmental and molecular mutagenesis 61(7):752–766. 10.1002/em.22401

Petr M, Pääbo S, Kelso J, et al. (2019) Limits of long-term selection against Neandertal introgression. Proceedings of the National Academy of Sciences of the United States of America 116(5):1639– 1644. 10.1073/pnas.1814338116

Pfennig A, Lachance J (2022) Hybrid fitness effects modify fixation probabilities of introgressed alleles. G3 Genes|Genomes|Genetics 12(7):jkac113. 10.1093/g3journal/jkac113

Prüfer K, Racimo F, Patterson N, et al. (2013) The complete genome sequence of a Neanderthal from the Altai Mountains. Nature 2013 505:7481 505(7481):43–49. 10.1038/nature12886

Prüfer K, De Filippo C, Grote S, et al. (2017) A high-coverage Neandertal genome from Vindija Cave in Croatia. Science 358(6363):655–658. 10.1126/science.aao1887

Quinlan AR, Hall IM (2010) BEDTools: a flexible suite of utilities for comparing genomic features. Bioinformatics 26(6):841–842. 10.1093/bioinformatics/btq033

Racimo F, Marnetto D, Huerta-Sánchez E (2017) Signatures of Archaic Adaptive Introgression in Present-Day Human Populations. Molecular Biology and Evolution 34(2):296–317. 10.1093/molbev/msw216

Reilly PF, Tjahjadi A, Miller SL, et al. (2022) The contribution of Neanderthal introgression to modern human traits. Current Biology 32(18):R970–R983. 10.1016/j.cub.2022.08.027

Remke M, Hielscher T, Korshunov A, et al. (2011) FSTL5 Is a Marker of Poor Prognosis in Non-WNT/Non-SHH Medulloblastoma. Journal of Clinical Oncology 29(29):3852–3861. 10.1200/JCO.2011.36.2798

Rimphanitchayakit V, Tassanakajon A (2010) Structure and function of invertebrate Kazal-type serine proteinase inhibitors. Developmental & Comparative Immunology 34(4):377–386. 10.1016/j.dci.2009.12.004

Sachdeva H, Barton NH (2018a) Introgression of a block of genome under infinitesimal selection. Genetics 209(4):1279–1303. 10.1534/genetics.118.301018

Sachdeva H, Barton NH (2018b) Replicability of introgression under linked, polygenic selection. Genetics 210(4):1411–1427. 10.1534/genetics.118.301429

Saldivar JC, Hamperl S, Bocek MJ, et al. (2018) An intrinsic S/G2 checkpoint enforced by ATR. Science 361(6404):806–810. 10.1126/science.aap9346

Sankararaman S, Mallick S, Dannemann M, et al. (2014) The genomic landscape of Neanderthal ancestry in present-day humans. Nature 2014 507:7492 507(7492):354–357. 10.1038/nature12961

Sankararaman S, Mallick S, Patterson N, et al. (2016) The Combined Landscape of Denisovan and Neanderthal Ancestry in Present-Day Humans. Current Biology 26(9):1241–1247. 10.1016/j.cub.2016.03.037

Sherman BT, Hao M, Qiu J, et al. (2022) DAVID: a web server for functional enrichment analysis and functional annotation of gene lists (2021 update). Nucleic Acids Research 50(W1):W216–W221. 10.1093/nar/gkac194

Siepel A, Bejerano G, Pedersen JS, et al. (2005) Evolutionarily conserved elements in vertebrate, insect, worm, and yeast genomes. Genome Research 15(8):1034–1050. 10.1101/ gr.3715005

Skov L, Coll Macià M, Sveinbjörnsson G, et al. (2020) The nature of Neanderthal introgression revealed by 27,566 Icelandic genomes. Nature 2020 582:7810 582(7810):78–83. 10.1038/s41586-020-2225-9

Stefansson H, Petursson H, Sigurdsson E, et al. (2002) Neuregulin 1 and Susceptibility to Schizophrenia. The American Journal of Human Genetics 71(4):877–892. 10.1086/342734

Steinrücken M, Spence JP, Kamm JA, et al. (2018) Model-based detection and analysis of introgressed Neanderthal ancestry in modern humans. Molecular Ecology 27(19):3873–3888. 10.1111/mec.14565

Szklarczyk D, Gable AL, Lyon D, et al. (2019) STRING v11: protein-protein association networks with increased coverage, supporting functional discovery in genome-wide experimental datasets. Nucleic acids research 47(D1):D607–D613. 10.1093/nar/gky1131

Sánchez-Quinto F, Lalueza-Fox C (2015) Almost 20 years of Neanderthal palaeogenetics: adaptation, admixture, diversity, demography and extinction. Philosophical Transactions of the Royal Society B: Biological Sciences 370(1660):20130374. 10.1098/rstb.2013.0374, publisher: Royal Society

Telis N, Aguilar R, Harris K (2020) Selection against archaic hominin genetic variation in regulatory regions. Nature Ecology & Evolution 2020 4:11 4(11):1558–1566. 10.1038/s41559-020-01284-0

Uecker H, Setter D, Hermisson J, et al. (2015) Adaptive gene introgression after secondary contact. Journal of Mathematical Biology 70:1523–1580. 10.1007/s00285-014-0802-y

All of Us Research Program Investigators, Denny J, Rutter J, et al. (2019) The “all of us” research program. New England Journal of Medicine 381(7):668–676. 10.1056/NEJMsr1809937

Vernot B, Akey JM (2014) Resurrecting surviving Neandertal lineages from modern human genomes. Science 343(6174):1017–1021. 10.1126/science.1245938

Vernot B, Tucci S, Kelso J, et al. (2016) Excavating Neandertal and Denisovan DNA from the genomes of Melanesian individuals. Science 352(6282):235–239. 10.1126/science.aad9416

Virtanen P, Gommers R, Oliphant TE, et al. (2020) SciPy 1.0: Fundamental Algorithms for Scientific Computing in Python. Nature Methods 17:261–272. 10.1038/s41592-019-0686-2

Walt Svd, Schönberger JL, Nunez-Iglesias J, et al. (2014) scikit-image: image processing in Python. PeerJ 2:e453. 10.7717/peerj.453

Wang RJ, Ané C, Payseur BA (2013) The evolution of hybrid incompatibilities along a phylogeny. Evolution 67(10):2905–2922. 10.1111/evo.12173

Wei X, Robles CR, Pazokitoroudi A, et al. (2023) The lingering effects of Neanderthal introgression on human complex traits. eLife 12:e80757. 10.7554/eLife.80757

Witt KE, Funk A, Añorve-Garibay V, et al. (2023) The Impact of Modern Admixture on Archaic Human Ancestry in Human Populations. Genome Biology and Evolution 15(5):evad066. 10.1093/gbe/evad066

Wohns AW, Wong Y, Jeffery B, et al. (2022) A unified genealogy of modern and ancient genomes. Science 375(6583):eabi8264. 10.1126/science.abi8264

Yates A, Beal K, Keenan S, et al. (2014) The Ensembl REST API: Ensembl Data for Any Language. Bioinformatics 31(1):143–145. 10.1093/bioinformatics/btu613

Zeberg H, Jakobsson M, Pääbo S (2024) The genetic changes that shaped Neandertals, Denisovans, and modern humans. Cell 187(5):1047–1058. 10.1016/j.cell.2023.12.029

Zhang D, Ma X, Sun W, et al. (2015) Down-regulated FSTL5 promotes cell proliferation and survival by affecting Wnt/*β*-catenin signaling in hepatocellular carcinoma. International Journal of Clinical and Experimental Pathology 8(3):3386–3394

Zhao H, Sun Z, Wang J, et al. (2013) CrossMap: a versatile tool for coordinate conversion between genome assemblies. Bioinformatics 30(7):1006–1007. 10.1093/bioinformatics/btt730

