## Supplemental Information for "The evolutionary fate of Neanderthal DNA in 30,780 admixed genomes with recent African-like ancestry"

#### Genome-wide trends from All of Us are replicated in 93 admixed individuals from 1KGP-ACB and 1KGP-ASW using different Neanderthal reference genomes

We replicated major genome-wide trends from All of Us in admixed populations from the 1000 genomes project (1KGP) phase 3, i.e., 1KGP-ACB and 1KGP-ASW (Auton et al., 2015), and each of the three available high-quality Neanderthal reference genomes (i.e., Altai, Vindija33.19, and Chagyrskaya) (Prüfer et al., 2013, 2017; Mafessoni et al., 2020). We first estimated genome-wide autosomal ancestry proportions for unrelated individuals from 1KGP-ACB and 1KGP-ASW to identify admixed individuals that conform our ancestry filters (i.e., at least 50% African-like, 10% European-like, and at most 5% East Asian/Native American-like ancestry). For this purpose, we constructed continental reference panels using 1KGP-AFR, -EUR, and -EAS populations as well as HGDP-Maya and HGDP-Pima populations (Auton et al., 2015; Bergström et al., 2020). Genotype calls for 1KGP-AFR, -EUR, and -EAS reference populations were lifted over from hg19 to hg38 using CrossMap v0.6.5 (Zhao et al., 2013) as described in the main text. We then merged 1KGP, HGDP-Maya, and HGDP-Pima data on the intersection of biallelic SNPs using plink2 (Chang et al., 2015). Using plink2, this set of shared biallelic SNPs was then LD pruned (*–indep-pairwise 200 kb 1 0.2*), and the top 20 principal components were computed on the LD pruned dataset (*–pca 20 approx*). Subsequently, we projected 1KGP-ACB and 1KGP-ASW individuals onto the reference PC space using the *–score* function in plink2. Finally, Rye v0.1 was used to determine ancestry proportions from the computed eigenvectors and eigenvalues for each of the granular reference populations (Conley et al., 2023), which were subsequently aggregated to continental ancestry proportions. We aggregated 1KGP-YRI, 1KGP-GWD, 1KGP-ESN, 1KGP-LWK, and 1KGP-MSL to African-like ancestry, 1KGP-CEU, 1KGP-FIN, 1KGP-GBR, 1KGP-TSI, and 1KGP-IBS to European-like ancestry, and 1KGP-CHB, 1KGP-CHS, 1KGP-JPT, 1KGP-CDX, 1KGP-KHV, HGDP-Maya, and HGDP-Pima to East Asian/Native American-like ancestry. The inclusion of HGDP-Maya and HGDP-Pima samples improved the painting of Native American-like ancestry, which is most similar to East Asian-like ancestry on a continental level.

We identified 93 admixed individuals with at least 50% African-like, 10% European-like, and at most 5% East Asian/Native American-like ancestry 1KGP-ACB and 1KGP-ASW. We analyzed this set of admixed individuals in the same way as the admixed individuals from All of Us, with the exception of not including additional reference individuals from All of Us (i.e., 563 African-like, 10,000 European-like, and 71 Native American-like individuals). However, we analyzed this test dataset using each of the three available high-quality Neanderthal reference genomes.

Similar to the patterns observed in the 30,780 admixed individuals from All of Us, we initially observed more Neanderthal ancestry than expected based on ancestry patterns (Figure S7A-F). However, this enrichment disappears after applying the African mask, i.e., removing any Neanderthal introgressed segment that overlaps with an introgressed segment in an African reference genome (Figure S7G -L), providing no evidence for strong polygenic selection of remaining Neanderthal ancestry on a genome level since admixture. Furthermore, our results are robust to the choice of Neanderthal reference genome as we observed qualitative similar patterns independent of which Neanderthal individual was used (Figure S7).

#### Accounting for genetic drift when probabilistically identifying 50 kb windows with significantly less or more Neanderthal ancestry is not well calibrated

Given our sample size of 30,780 admixed individuals, we have statistical power to detect small deviations in the expected introgression frequencies that are conceivably explained by genetic drift using Equation 3 in the main text (see example below). For this reason, we accounted for the expected variance in introgression frequencies for each window $i$ after 15 generations of drift. The expected variance in allele frequencies after $t$ generations of drift in haploid genomes is given by:

$$V = pq(1 - exp(-\frac{t}{N_e})) \tag{S1}$$

where $p$ and $q$ are the starting introgression frequencies, and $N_e$ is the effective population size (Charlesworth and Charlesworth, 2010). We can use Equation S1 to calculate expected variances in introgression frequencies because we consider short regions (i.e., 50 kb windows) and therefore can treat introgression frequencies like alleles frequencies and because we deal with pseudo-haploid genomes since IBDmix does not provide phase information.

To illustrate that we have statistical power to detect deviations within one standard deviation of expected introgression frequencies after 15 generations of drift, we consider the following example. Assume an introgression frequency of 0.004 in the admixed individuals, i.e., an introgression frequency of 0.02 in European populations and a European-like admixture fraction of 0.2. In this case, we have approximately 80% power to detect an absolute depletion or enrichment of 0.001 at a significance level of 0.05. However, in this example and assuming a $N_e$ of 30,000, the standard deviation in introgression frequencies after 15 generations of drift is 0.0014 (Equation S1). Thus, we have 80% power to detect deviations within one standard deviation of expected introgression frequencies after 15 generations of genetic drift, and many of the depletion/enrichments identified using Equation 3 in the main text would be false positives.

To identify 50 kb windows with significantly less or more Neanderthal ancestry than expected based on local ancestry patterns and introgression frequencies in the source populations, we only considered windows with expected introgression frequency greater than zero, less than 50% masked sites, intermediate recombination rate (i.e., $\geq$0.65 cm/Mb and $\leq$1.52 cM/Mb), and that have at least 50% African-like, at least 10% European-like, and less than 5% East Asian/Native American-like ancestry. We then calculated the probability of observing a given enrichment depletion using Equation 3 in the main text, performed a Bonferroni correction, and required significant depletions and enrichments to be stronger than could be expected due to genetic drift, i.e., outside of the Bonferroni corrected 95% confidence interval (where $n$ is the number tested windows) of the expected introgression frequencies after 15 generations of drift for each window (Equation S1).

We identified 646 outlier windows (Figure S8A) that form six and 124 independent genomic regions that are depleted and enriched for Neanderthal ancestry, respectively. However, neutral simulations suggest that many of these genomic regions are false positives, and in particular, enriched regions appear to be false positives. In neutral coalescence simulations, we identified between 0 to 22 windows with significantly less Neanderthal ancestry than expected and 57 to 108 windows with significantly more Neanderthal ancestry than expected, respectively (Figure S9). For this reason, we ultimately conditioned the identification of regions with significantly less or more Neanderthal ancestry than expected on the simulated joint spectrum of expected and observed introgression frequencies as described in the main text.

#### Dating of genetic variants

For the sake of computational tractability, we only estimated allele ages using phased genotype data of continental 1KGP reference populations (Auton et al., 2015), i.e., 1KGP-AFR, 1KGP-EUR, and 1KGP-EAS, and the four archaic reference genomes (i.e., Altai, Vindija33.19, and Chagyrskaya Neanderthals and Denisovan) (Prüfer et al., 2013, 2017; Mafessoni et al., 2020; Meyer et al., 2012), considering only biallelic sites for which the ancestral allele status was available. Ancestral allele states were inferred from 10 primates' EPO alignments using the ENSEMBL Compara Perl API (Yates et al., 2014; Herrero et al., 2016; Cunningham et al., 2021). For each chromosome arm, an initial tree sequence was inferred using tsinfer v0.3.0 (Kelleher et al., 2019) with HapMap recombination maps for hg19 (Frazer et al., 2007). Subsequently, allele ages were estimated using tsdate v0.1.5 (Wohns et al., 2022), assuming an effective population size of 10,000 and a mutation rate of $1.25 \times 10^{-8}$ per base pair per generation. Allele ages were constrained using radiocarbon dates of the archaic reference genomes. For derived alleles present in an archaic individual, the lower bound was set to the radiocarbon date of the oldest archaic reference genomes that carried the derived allele, i.e., 110 kya for the Altai Neanderthal, 50 kya for the Vindija33.19 Neanderthal, 80 kya for the Chagyrskaya Neanderthal, and 63.9 kya for the Denisovan individual, assuming a generation time of 29 years. With these constraints, the tree sequence was then re-inferred. Estimated allele ages were then converted to years by again assuming a generation time of 29 years. Finally, we lifted over coordinates from hg19 to hg38 using CrossMap v0.6.5 (Zhao et al., 2013) and masked previously inferred CpG sites (see Materials and Methods in the main text). We compared estimated allele ages of Neanderthal-derived variants in novel desert-like regions, previously known deserts (Vernot et al., 2016; Chen et al., 2020), and the genomic background. We defined Neanderthal-derived variants as variants that are present in one or more Neanderthal individuals while not being found in the Denisovan individual or African individuals. We used a Mann-Whitney U test as implemented in scipy v1.10.1 to assess statistical significance (Virtanen et al., 2020).

#### Supplemental Figures

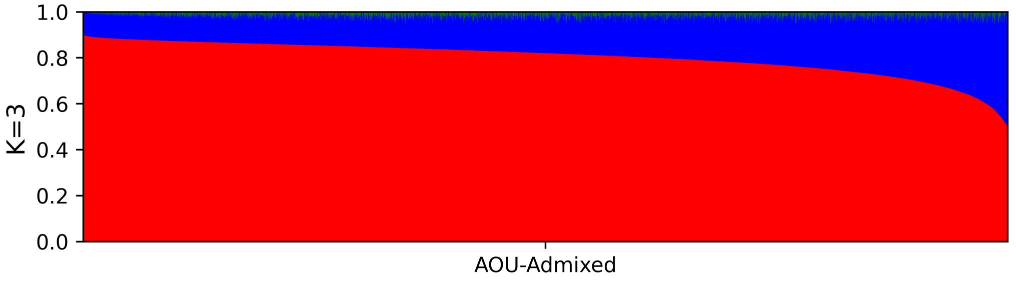


**Fig. S1** Inferred ancestry proportions of 30,780 admixed individuals with at least 50% recent African-like ancestry (red), 10% European-like ancestry (blue), and less than 5% East Asian/Native American-like ancestry (green) from All of Us. Due to the difficulty of painting Native American-like ancestry, we summed inferred East Asian-like and Native American-like ancestry proportions. Ancestry proportions were previously estimated using Rye (Conley et al., 2023; All of Us Research Program Investigators et al., 2019; Bick et al., 2024). Because Rye is a supervised method (Conley et al., 2023), ancestry proportions are estimated using a fixed number of components. For this reason, no other values for $K$ were explored. Related to Materials and Methods.

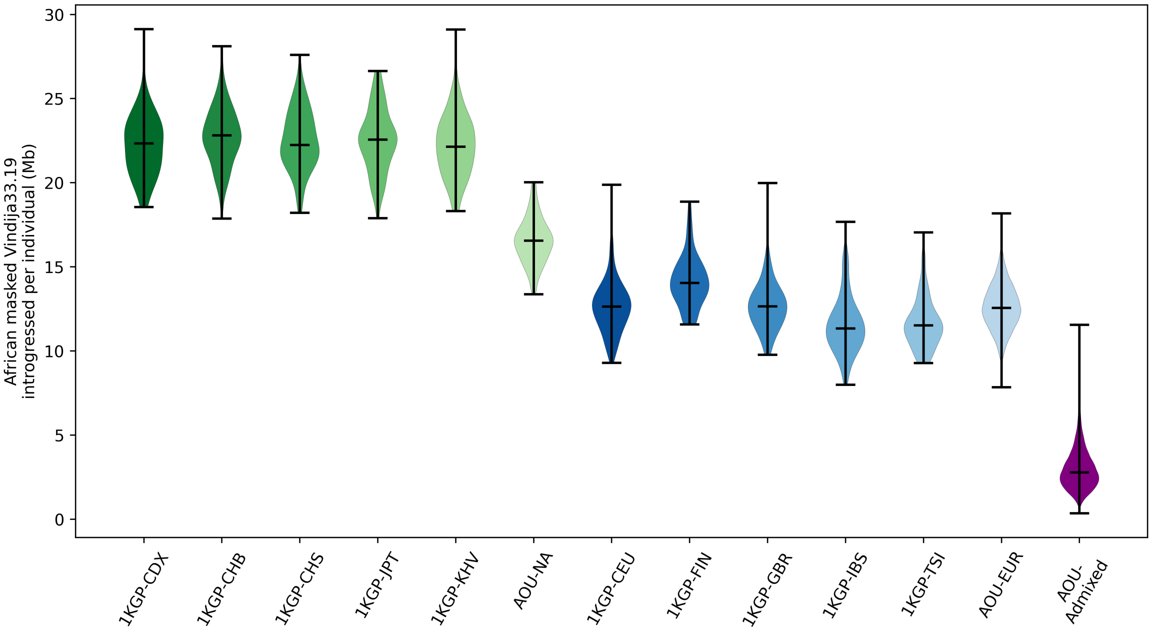


**Fig. S2** Inferred amounts of Neanderthal ancestry in Mb per individual in global populations after applying the African mask, i.e., removing any introgressed segment overlapping with an introgressed segment in an African reference genome by at least 1 bp. The masking removes more introgressed sequence from European reference genomes than East Asian/Native American reference genomes because most true Neanderthal sequence in African genomes comes from European back migration (Chen et al., 2020). European and East Asian/Native American reference genomes contain, on average, 12.6 Mb and 21.7 Mb introgressed Neanderthal sequence, respectively, while admixed genomes contain, on average, 3 Mb of unmasked Neanderthal introgressed sequence. For this figure, the high-quality Vindija33.19 Neanderthal genome was used as a Neanderthal reference. Related to Figure 2A and Materials and Methods.

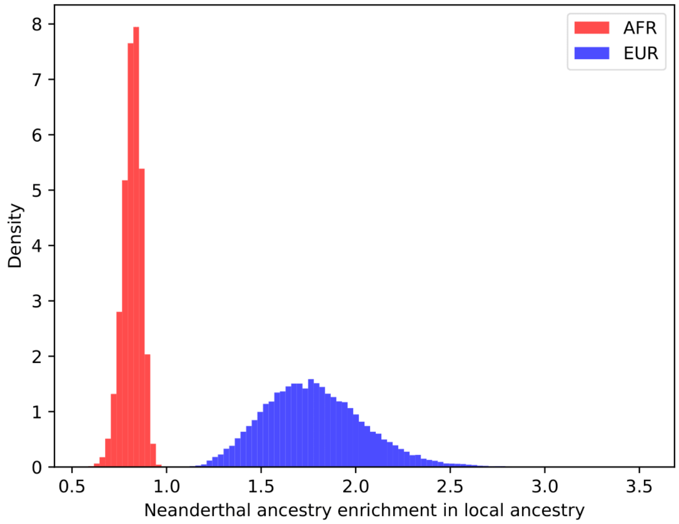


**Fig. S3** Overlap of Neanderthal introgressed segments with local ancestry tracks. Neanderthal introgressed segments overlap with tracks of local European-like ancestry more frequently than expected by chance. The enrichment was calculated by computing the proportion of Neanderthal segments overlapping with tracks of a given local ancestry in either phase and then dividing by the genome-wide ancestry proportion. For this figure, the high-quality Vindija33.19 Neanderthal genome was used as a Neanderthal reference. Related to Figure 2B-D.

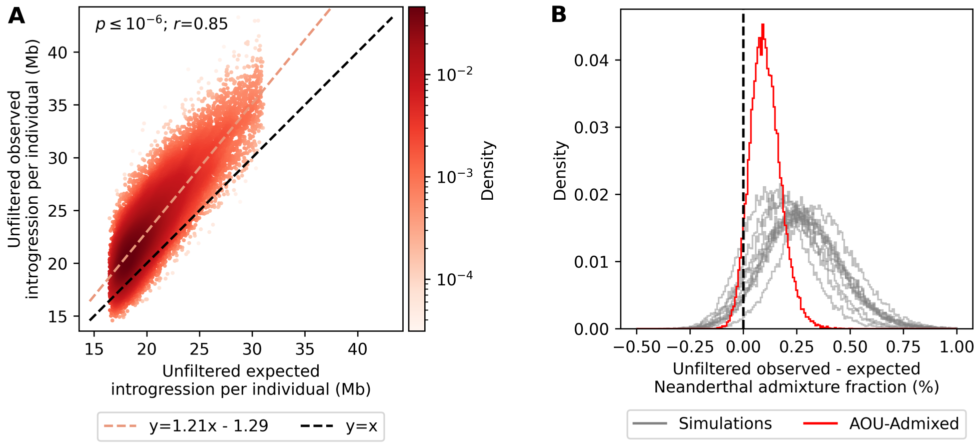


**Fig. S4** Admixed genomes contain more Neanderthal ancestry than expected when considering all predicted introgressed segments. **A)** Comparison of expected and observed amounts of Neanderthal introgressed sequence per individual in Mb. The slope of the regression line is significantly greater than one (m=1.21 95% CI: 1.20-1.22), and the intercept is less than zero (b=-1.29 95% CI:-1.46 - -1.12). The p-value ($p$) and Pearson's correlation coefficient ($r$) of the regression line are given in the panel. **B)** Differences in expected and observed Neanderthal admixture fractions are shifted to values greater than zero for empirical data (red) and data from neutral coalescence simulations (gray). The mean difference in the Neanderthal admixture fraction in the empirical data is 3 Mb (0.104% of the entire genome). For this figure, the high-quality Vindija33.19 Neanderthal genome was used as a Neanderthal reference. Related to Figure 3.

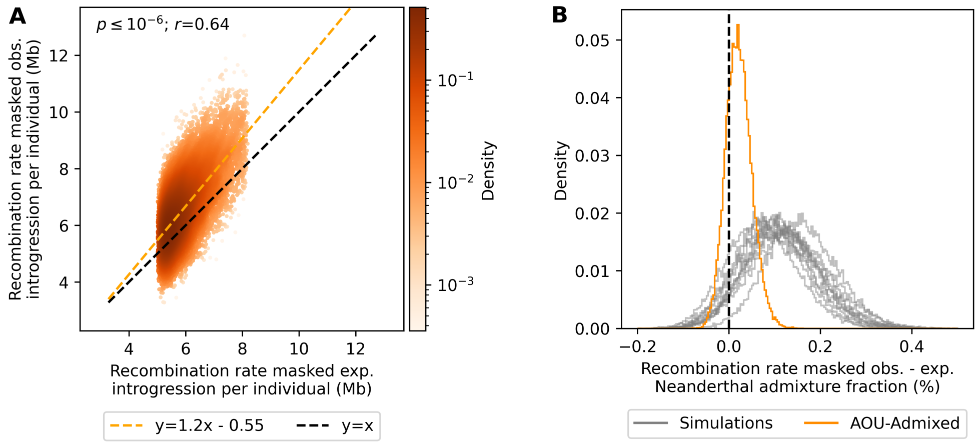


**Fig. S5** Admixed genomes contain more Neanderthal ancestry than expected after removing segments in low and high recombination rate regions. Predicted Neanderthal introgressed segments overlapping with 300 kb windows in the bottom and top 33rd percentiles in terms of recombination rate were removed (i.e., $< 0.65$ cm/Mb or $> 1.52$ cM/Mb). **A)** Comparison of expected and observed amounts of Neanderthal introgressed sequence per individual in Mb. The slope of the regression line remains significantly greater than one (m=1.20 95% CI:1.19-1.22), and the intercept is less than zero (b=-0.55 95% CI:-0.64 - -0.45). The p-value ($p$) and Pearson's correlation coefficient ($r$) of the regression line are given in the panel. **B)** Differences in expected and observed Neanderthal admixture fractions are shifted to values greater than zero for empirical data (orange) and data from neutral coalescence simulations (gray). The mean difference in the Neanderthal admixture fraction in the empirical data is 0.63 Mb (0.022% of the entire genome). For this figure, the high-quality Vindija33.19 Neanderthal genome was used as a Neanderthal reference. Related to Figure 3.

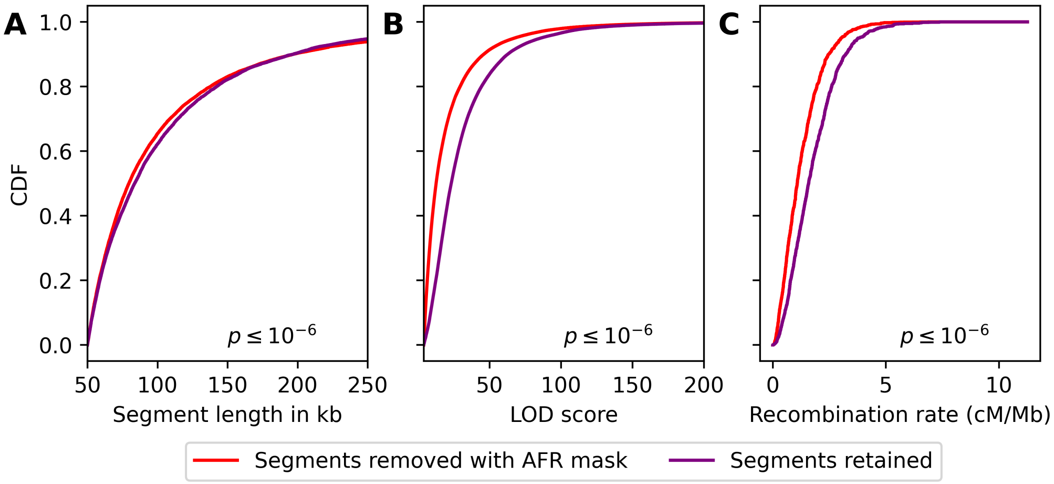


**Fig. S6** Comparison of introgressed segments that are removed (red) or retained (purple) with the African mask, i.e., segments that overlap with a segment in African reference genomes, in terms of length **(A)**, LOD scores **(B)**, and recombination **(C)**. Segments that are removed by the African mask are significantly shorter, have lower LOD scores, and are in lower recombination rate regions than segments that were retained, indicating that they are more likely false positive predictions by IBDmix. Statistical significance was assessed using a Mann-Whitney U test, and corresponding p-values are provided in each panel. For this figure, the high-quality Vindija33.19 Neanderthal genome was used as a Neanderthal reference. Related to Figure 3 and Materials and Methods.

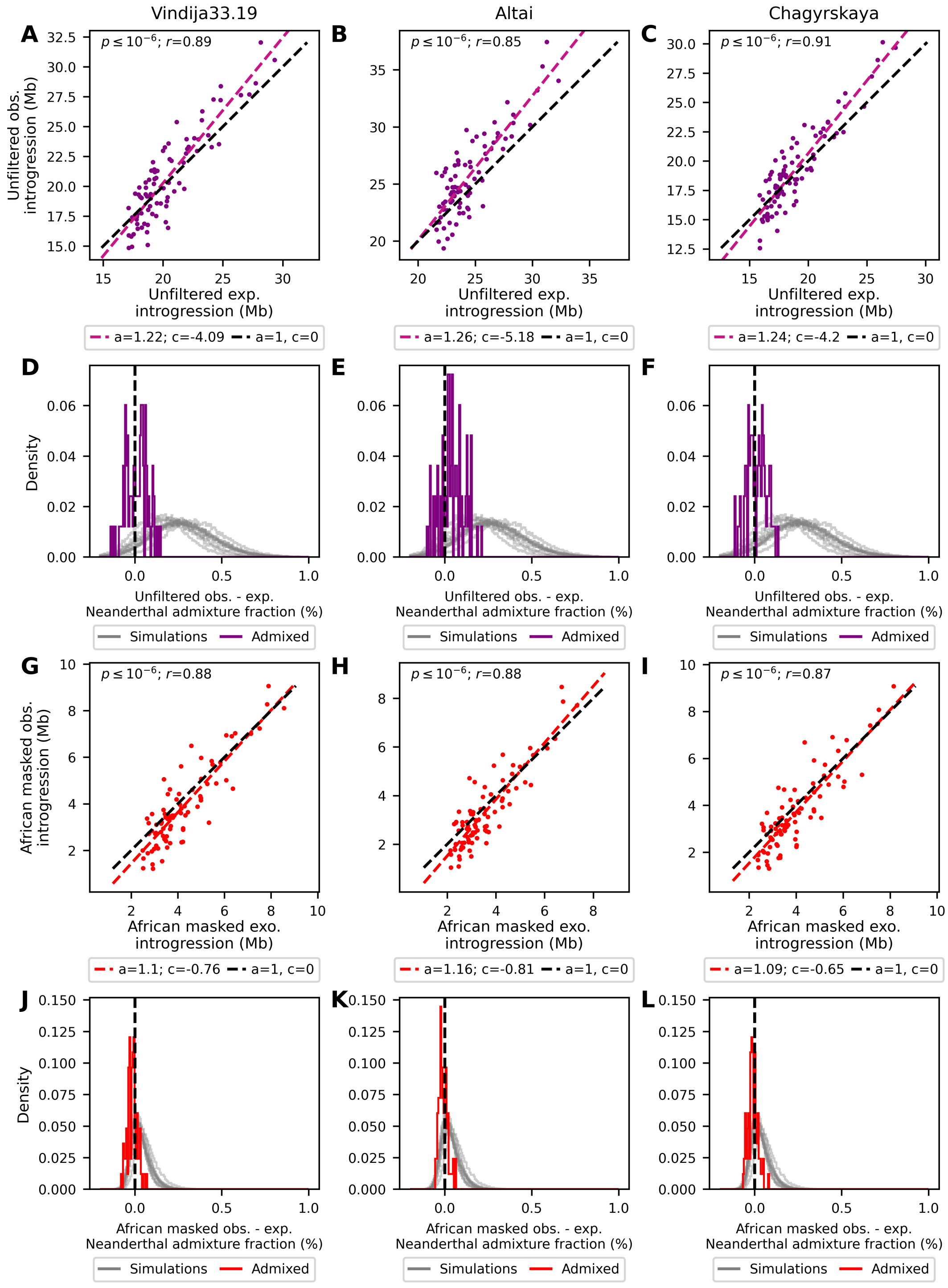


**Fig. S7** Comparison of the expected versus observed amounts of Neanderthal ancestry in 93 admixed genomes with predominantly African-like and European-like ancestry from 1KGP-ACB and 1KGP-ASW when using all available Neanderthal reference genomes. **A-C)** and **D-F)** show the comparison when all segments are considered, while **G-I)** and **J-L)** show the comparison after masking introgressed segments also found in African reference genomes. The left column shows the results using the Vindija33.19 Neanderthal reference genome, the middle column using the Altai Neanderthal reference genome, and the right column using the Chagyrskaya reference genome. All Neanderthal reference genomes yield qualitatively similar results to those observed in 30,780 admixed genomes from All of Us. The p-value ($p$) and Pearson's correlation coefficients ($r$) of linear regressions are indicated in the respective panels. Related to Figure 3 and Materials and Methods.

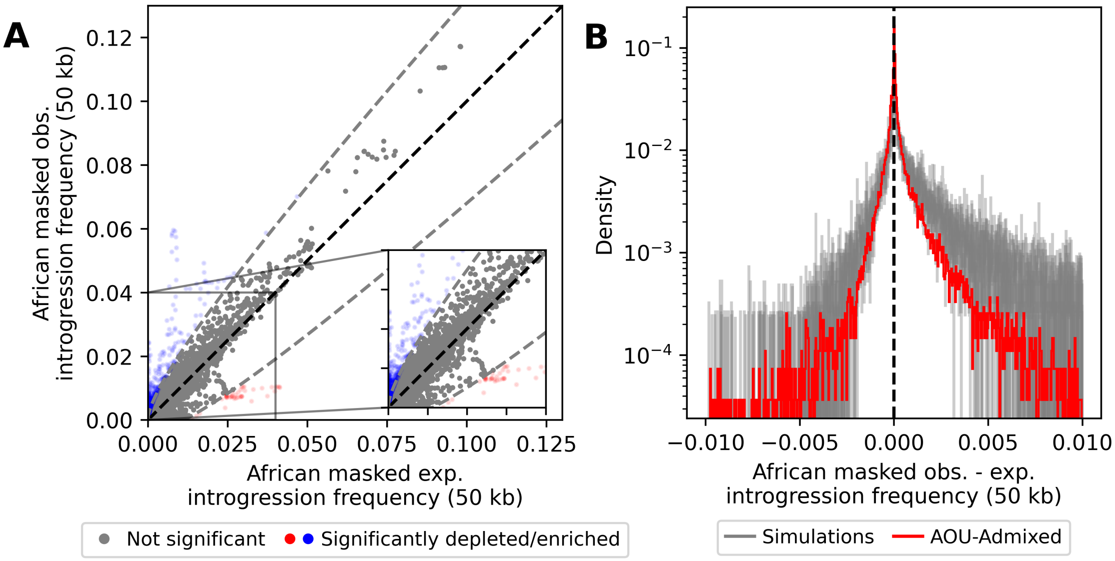


**Fig. S8** Expected versus observed Neanderthal introgression frequencies in 50 kb windows in the 30,780 admixed individuals from All of Us after applying the African mask. **A)** Many windows are found to have significantly less (red) and more (blue) Neanderthal ancestry than expected on a Bonferroni-corrected significance level of 0.05 and after accounting for genetic drift. The dashed grey lines correspond to the Bonferroni corrected 95% confidence interval of expected frequencies after 15 generations of drift (Equation S1). **B)** Differences between expected and observed introgression frequencies in 50 kb windows. The modeling of expected introgression frequencies in 50 kb windows is generally unbiased, as the differences are centered around 0. Only windows with an expected introgression frequency greater than 0, less than 50% masked sites, intermediate recombination rate (i.e., $\geq$0.65 cm/Mb and $\leq$1.52 cM/Mb), and that have at least 50% African-like, at least 10% European-like, and less than 5% East Asian/Native American-like ancestry were included in the analyses. For this figure, the high-quality Vindija33.19 Neanderthal genome was used as a Neanderthal reference. Related to Figure 4 and Materials and Methods.

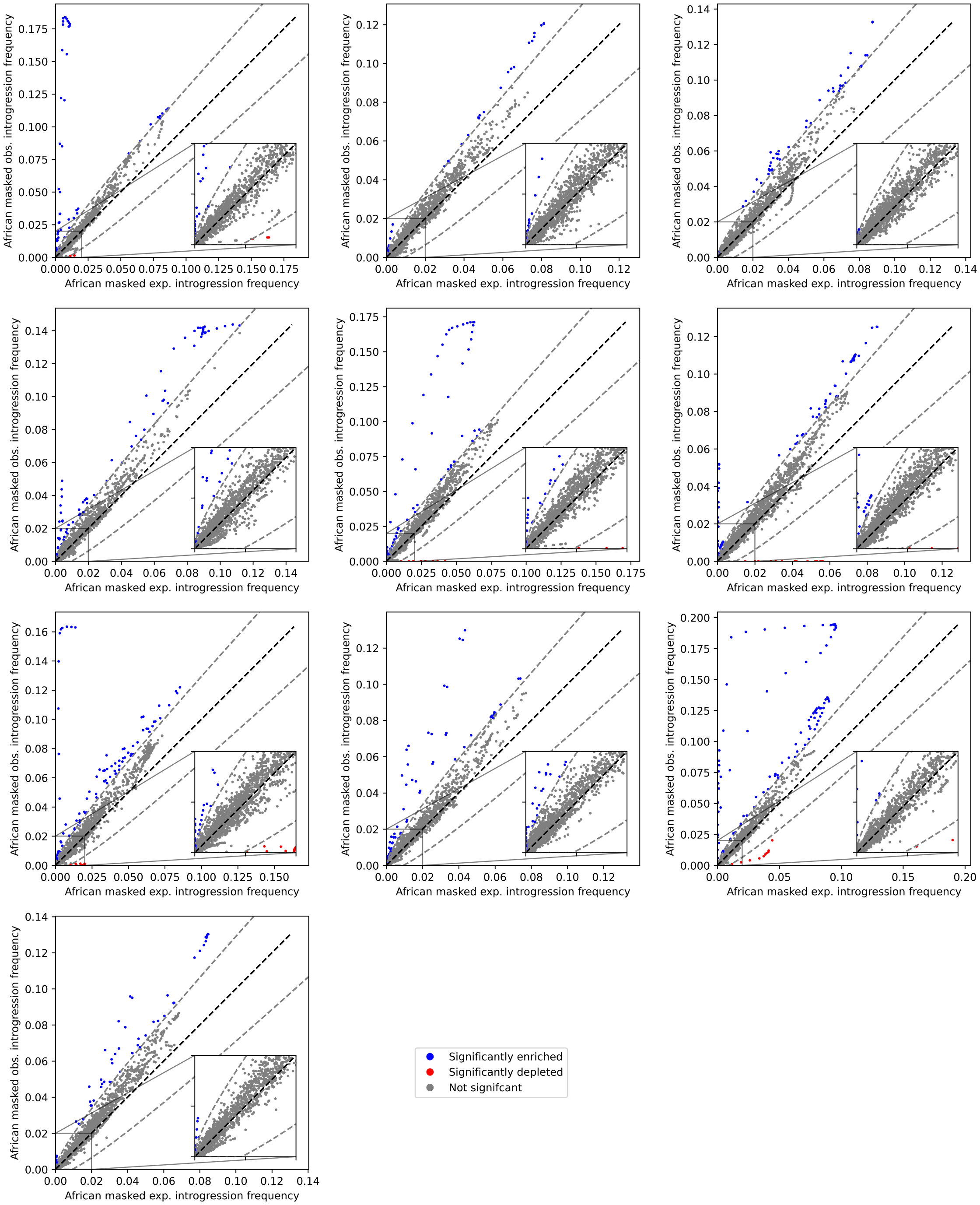


**Fig. S9** Expected vs. observed Neanderthal introgression frequencies in overlapping 50 kb (10 kb step size) in African masked call set of Neanderthal segments for each of the ten neutral simulation replicates. Windows with significantly less or more Neanderthal ancestry than expected on a Bonferroni-corrected significance level of 0.05 and after accounting for the variance in expected introgression frequencies due to 15 generations of drift are shown in red and blue, respectively. The dashed grey lines correspond to the Bonferroni corrected 95% confidence interval of expected frequencies after 15 generations of drift (Equation S1). Given the number of significant outlier windows, these simulations indicate that our modeling approach (Equations 2-S1) is not well calibrated. Only windows with an expected introgression frequency greater than 0, less than 50% masked sites, intermediate recombination rate (i.e., $\geq$0.65 cm/Mb and $\leq$1.52 cM/Mb), and that have at least 50% African-like, at least 10% European-like, and less than 5% East Asian/Native American-like ancestry were included in the analyses. Related to Figure 4 and Materials and Methods.

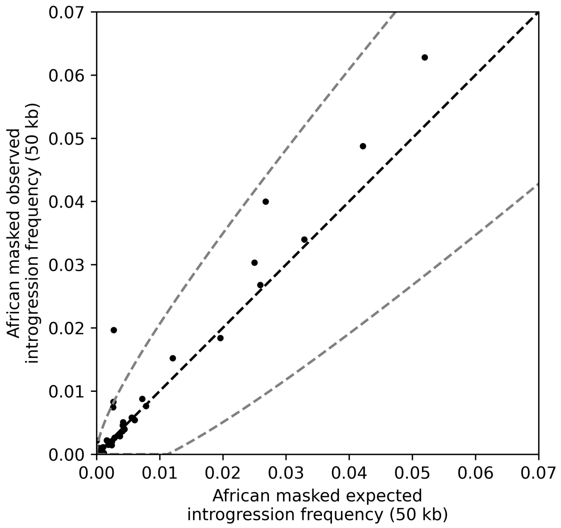


**Fig. S10** Comparison of expected and observed introgression frequencies for 93 candidate regions for adaptive Neanderthal introgression in European populations identified by Racimo et al. (2017). These regions generally have introgression frequencies as expected based on local ancestry patterns and introgression frequencies in the admixing reference populations. Three regions have a higher Neanderthal introgression frequency than could be expected after 15 generations of drift: chr5:168652996-168692995, chr9:16800003-16840002, and chr18:53993631-54033630. For this figure, the high-quality Vindija33.19 Neanderthal genome was used as a Neanderthal reference, and genomic positions are given relative to hg38. Related to Figure 4 and Materials and Methods.

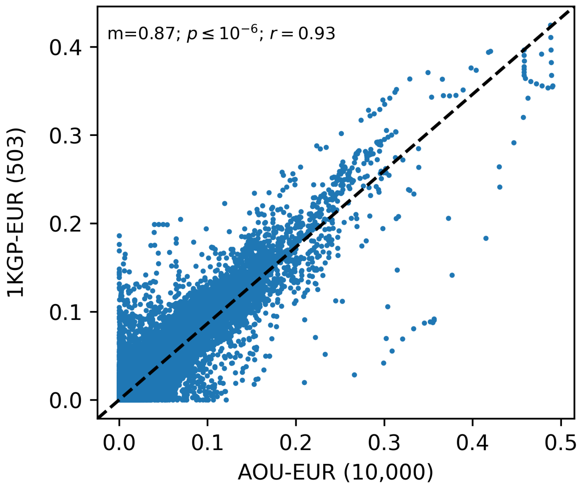


**Fig. S11** Comparison of calculated Neanderthal introgression frequencies in 50 kb windows in 503 and 10,000 European-like individuals from 1KGP and All of Us, respectively. The frequencies are highly correlated with a $r = 0.93$. Introgression frequencies were calculated based on the African masked call set of introgressed segments using the high-quality Vindija33.19 Neanderthal genome. The slope ($m$), p-value ($p$), and Pearson's correlation coefficients ($r$) of a linear regression are indicated in the Figure. Related to Materials and Methods.

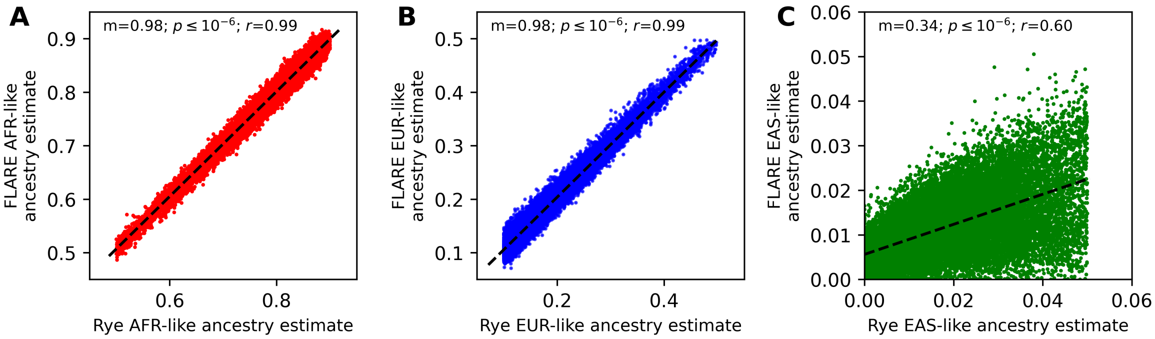


**Fig. S12** Comparison of previously inferred ancestry proportions and global ancestry estimates by FLARE for the 30,780 admixed individuals with predominantly African-like and European-like ancestry from All of Us. Previously inferred global ancestry proportions using Rye (All of Us Research Program Investigators et al., 2019; Conley et al., 2023; Bick et al., 2024) are highly correlated with **(A)** African-like, **(B)** European-like, and **(C)** East Asian/Native American-like ancestry estimates by FLARE. The slope ($m$), p-value ($p$), and Pearson's correlation coefficients ($r$) of linear regressions are indicated in the respective panel. Related to Materials and Methods.

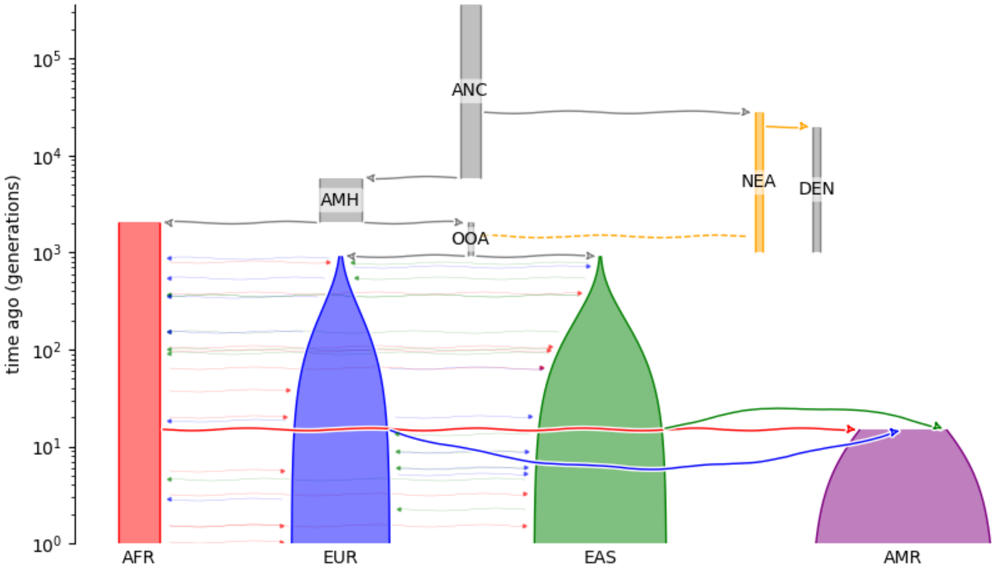


**Fig. S13** Visualization of the simulated demographic model. The three populations out-of-Africa model by Gravel et al. (2011) was extended to include archaic introgression and recent admixture in the Americas following Browning et al. (2018). The demographic model was visualized using demesdraw (Gower, 2023). Related to Materials and Methods. Related to Materials and Methods.

#### Supplemental Tables

**Table S1** Regions containing significantly less or more Neanderthal ancestry than expected. Protein-coding genes within 500 kb downstream and upstream of the regions are reported. All coordinates are relative to hg38. Expected and observed introgression frequencies were calculated based on the African masked call set. Related to Figure 5.

| **Genomic region** | **African masked expected Neanderthal introgression frequency** | **African masked observed Neanderthal introgression frequency** | **Genes (GENCODE v46)** |
|---|---|---|---|
| chr2:67,300,000-67,450,000 | 0.024644 | 0.007638 | *ETAA1* |
| chr4:160,900,000-161,100,000 | 0.028262 | 0.008363 | *FSTL5* |
| chr8:32,470,000-32,550,000 | 0.025480 | 0.007232 | *NRG1, ENSG00000286131* |
| chr14:44,170,000-44,340,000 | 0.035520 | 0.008944 | *FSCB* |
| chr4:100,840,000-100,900,000 | 0.014907 | 0.042528 | *EMCN, PPP3CA* |
| chr5:147,670,000-147,800,000 | 0.008421 | 0.052274 | *C5orf46, DPYSL3, JAKMIP2, MARCOL, SCGB3A2, STK32A, SPINK1, SPINK5, SPINK6, SPINK13, SPINK14* |
| chr6:76,660,000-76,800,000 | 0.016505 | 0.044031 | - |

**Table S2** Genomic coordinates of novel desert-like regions identified in this study and previously known introgression deserts from Vernot et al. (2016) and Chen et al. (2020). The novel desert-like region on chromosome 7 overlaps with a previously known desert and, for this reason, was excluded from subsequent analyses. Retained novel desert-like regions are shown in boldface. All coordinates are relative to hg38. Coordinates of previously known deserts from Vernot et al. (2016) and Chen et al. (2020) were lifted over from hg19 to hg38 using CrossMap v0.6.5 (Zhao et al., 2013) and merged using bedtools v2.30.0 (Quinlan and Hall, 2010). Expected and observed introgression frequencies were calculated based on the African masked call set. Related to Figure 6.

| **Chromosome** | **Start** | **End** | **African masked exp. Neanderthal introgression frequency** | **African masked obs. Neanderthal introgression frequency** | **Source** |
|---|---|---|---|---|---|
| 1 | 101,734,444 | 120,057,386 | $2.294 \times 10^{-4}$ | $1.352 \times 10^{-4}$ | Vernot et al. (2016), Chen et al. (2020) |
| **2** | **18,900,000** | **28,300,000** | $1.123 \times 10^{-3}$ | $2.078 \times 10^{-4}$ | **This study** |
| 2 | 200,235,277 | 210,635,276 | $3.937 \times 10^{-4}$ | $5.492 \times 10^{-4}$ | Vernot et al. (2016) |
| 3 | 74,050,849 | 90,450,850 | $2.858 \times 10^{-4}$ | $1.757 \times 10^{-4}$ | Vernot et al. (2016), Chen et al. (2020) |
| 7 | 106,559,554 | 125,059,946 | $8.298 \times 10^{-4}$ | $9.747 \times 10^{-5}$ | Vernot et al. (2016), Chen et al. (2020) |
| 7 | 110700000 | 118800000 | $6.881 \times 10^{-4}$ | $2.475 \times 10^{-4}$ | This study |
| 8 | 48,487,440 | 65,587,765 | $3.068 \times 10^{-4}$ | $2.594 \times 10^{-4}$ | Vernot et al. (2016), Chen et al. (2020) |
| **10** | **48,300,000** | **59,800,000** | $1.067 \times 10^{-3}$ | $1.735 \times 10^{-4}$ | **This study** |
| **17** | **49,500,000** | **64,600,000** | $2.222 \times 10^{-3}$ | $6.401 \times 10^{-4}$ | **This study** |
| 18 | 27,420,036 | 44,220,035 | $2.344 \times 10^{-4}$ | $2.710 \times 10^{-4}$ | Vernot et al. (2016) |

**Table S3** Biological processes with overrepresentation of genes among 243 genes overlapping with novel introgression desert-like regions compared against the human genome background (19,256 genes) reported by DAVID (Sherman et al., 2022). Related to Figure 6.

| GO Biological Process Direct | # Background genes | # genes in list | Enrichment | P-value | FDR |
|---|---|---|---|---|---|
| Reproductive process | 1611 | 33 | 1.6 | $6.6 \times 10^{-3}$ | $5.2 \times 10^{-2}$ |
| Reproduction | 1616 | 33 | 1.6 | $6.9 \times 10^{-3}$ | $5.2 \times 10^{-2}$ |
| Localization | 6551 | 102 | 1.2 | $7.1 \times 10^{-3}$ | $5.2 \times 10^{-2}$ |
| Single-organism process | 14663 | 200 | 1.1 | $1.3 \times 10^{-2}$ | $7.4 \times 10^{-2}$ |
| Metabolic process | 12586 | 172 | 1.1 | $4.6 \times 10^{-2}$ | $2.0 \times 10^{-1}$ |
| Biological regulation | 13158 | 177 | 1.1 | $7.8 \times 10^{-2}$ | $2.4 \times 10^{-1}$ |
| Cellular process | 17578 | 228 | 1.0 | $9.6 \times 10^{-2}$ | $2.4 \times 10^{-1}$ |

### References

Auton A, Abecasis GR, Altshuler DM, et al. (2015) A global reference for human genetic variation. Nature 526(7571):68–74. https://doi.org/10.1038/nature15393

Bergström A, McCarthy SA, Hui R, et al. (2020) Insights into human genetic variation and population history from 929 diverse genomes. Science 367(6484):eaay5012. https://doi.org/10.1126/science.aay5012

Bick AG, Metcalf GA, Mayo KR, et al. (2024) Genomic data in the All of Us Research Program. Nature pp 1–7. https://doi.org/10.1038/s41586-023-06957-x

Browning SR, Browning BL, Daviglus ML, et al. (2018) Ancestry-specific recent effective population size in the Americas. PLOS Genetics 14(5):1–22. https://doi.org/10.1371/journal.pgen.1007385

Chang CC, Chow CC, Tellier LCAM, et al. (2015) Second-generation PLINK: rising to the challenge of larger and richer datasets. GigaScience 4(1). https://doi.org/10.1186/s13742-015-0047-8

Charlesworth B, Charlesworth D (2010) Elements of Evolutionary Genetics. W. H. Freeman

Chen L, Wolf AB, Fu W, et al. (2020) Identifying and Interpreting Apparent Neanderthal Ancestry in African Individuals. Cell 180(4):677–687.e16. https://doi.org/10.1016/j.cell.2020.01.012

Conley AB, Rishishwar L, Ahmad M, et al. (2023) Rye: genetic ancestry inference at biobank scale. Nucleic Acids Research https://doi.org/10.1093/nar/gkad149

Cunningham F, Allen JE, Allen J, et al. (2021) Ensembl 2022. Nucleic Acids Research 50(D1):D988–D995. https://doi.org/10.1093/nar/gkab1049

Frazer KA, Ballinger DG, Cox DR, et al. (2007) A second generation human haplotype map of over 3.1 million SNPs. Nature 449(7164):851–861. https://doi.org/10.1038/nature06258

Gower G (2023) Demesdraw. Available from: https://github.com/grahamgower/demesdraw

Gravel S, Henn BM, Gutenkunst RN, et al. (2011) Demographic history and rare allele sharing among human populations. Proceedings of the National Academy of Sciences of the United States of America 108(29):11983–11988. https://doi.org/10.1073/pnas.1019276108

Herrero J, Muffato M, Beal K, et al. (2016) Ensembl comparative genomics resources. Database 2016. https://doi.org/10.1093/database/bav096

Kelleher J, Wong Y, Wohns AW, et al. (2019) Inferring whole-genome histories in large population datasets. Nature Genetics 51(9):1330–1338. https://doi.org/10.1038/s41588-019-0483-y

Mafessoni F, Grote S, de Filippo C, et al. (2020) A high-coverage neandertal genome from chagyrskaya cave. Proceedings of the National Academy of Sciences 117(26):15132–15136. https://doi.org/10.1073/pnas.2004944117

Meyer M, Kircher M, Gansauge MT, et al. (2012) A high-coverage genome sequence from an archaic denisovan individual. Science 338(6104):222–226. https://doi.org/10.1126/science.1224344

Prüfer K, Racimo F, Patterson N, et al. (2013) The complete genome sequence of a Neanderthal from the Altai Mountains. Nature 2013 505:7481 505(7481):43–49. https://doi.org/10.1038/nature12886

Prüfer K, De Filippo C, Grote S, et al. (2017) A high-coverage Neandertal genome from Vindija Cave in Croatia. Science 358(6363):655–658. https://doi.org/10.1126/science.aao1887

Quinlan AR, Hall IM (2010) BEDTools: a flexible suite of utilities for comparing genomic features. Bioinformatics 26(6):841–842. https://doi.org/10.1093/bioinformatics/btq033

Racimo F, Marnetto D, Huerta-Sánchez E (2017) Signatures of Archaic Adaptive Introgression in Present-Day Human Populations. Molecular Biology and Evolution 34(2):296–317. https://doi.org/10.1093/molbev/msw216

Sherman BT, Hao M, Qiu J, et al. (2022) DAVID: a web server for functional enrichment analysis and functional annotation of gene lists (2021 update). Nucleic Acids Research 50(W1):W216–W221. https://doi.org/10.1093/nar/gkac194

All of Us Research Program Investigators , Denny J, Rutter J, et al. (2019) The "all of us" research program. New England Journal of Medicine 381(7):668–676. https://doi.org/10.1056/NEJMsr1809937

Vernot B, Tucci S, Kelso J, et al. (2016) Excavating Neandertal and Denisovan DNA from the genomes of Melanesian individuals. Science 352(6282):235–239. https://doi.org/10.1126/science.

aad9416

Virtanen P, Gommers R, Oliphant TE, et al. (2020) SciPy 1.0: Fundamental Algorithms for Scientific Computing in Python. Nature Methods 17:261–272. https://doi.org/10.1038/s41592-019-0686-2

Wohns AW, Wong Y, Jeffery B, et al. (2022) A unified genealogy of modern and ancient genomes. Science 375(6583):eabi8264. https://doi.org/10.1126/science.abi8264

Yates A, Beal K, Keenan S, et al. (2014) The Ensembl REST API: Ensembl Data for Any Language. Bioinformatics 31(1):143–145. https://doi.org/10.1093/bioinformatics/btu613

Zhao H, Sun Z, Wang J, et al. (2013) CrossMap: a versatile tool for coordinate conversion between genome assemblies. Bioinformatics 30(7):1006–1007. https://doi.org/10.1093/bioinformatics/btt730
